# Programmable gold nanoparticle conjugates enable precise AMPA receptor localization at brain synapses by cryo-ET

**DOI:** 10.64898/2026.09.18.752772

**Authors:** Cathy J Spangler, Gunasekaran Dhandapani, Aya Matsui, Elizaveta V Shiltseva, Jay Nix, Jonathan N Pruneda, Johannes Elferich, Eric Gouaux

## Abstract

Cryo-electron tomography (cryo-ET) is a powerful approach to image cells, organelles, viruses and molecular assemblies. However, resolving smaller complexes and molecules is challenging and presents a roadblock to the broad application of cryo-ET. Gold nanoparticles (AuNPs) have been used to localize otherwise unidentifiable molecules and complexes in tomograms but are typically tethered to the target of interest via a flexible linker. As a result, the position of the AuNP does not directly define the location of the target. Here we attach AuNPs to Fab antibody fragments via cysteine residues on defined elements of secondary structure and show that the AuNPs are rigidly bound to the Fab. We use these Fab-AuNP conjugates to locate AMPA receptors and probe their conformation upon desensitization with sub nanometer resolution in tomograms derived from brain slice lamella. By harnessing small, homogeneous AuNPs with precise conjugation to a Fab, we have developed reagents that enable direct investigation of conformational states of previously unidentifiable biomolecules and complexes in tomograms.

## Main

Cryogenic electron microscopy^1^ (cryo-EM) has emerged as the preeminent experimental strategy to elucidate biological molecular structure and cellular architecture, with single particle methods^2^ enabling atomic resolution and tomographic approaches^3^ reaching sub nanometer resolution. Whereas single particle approaches are now able to determine the structures of molecules under 50 kilodaltons (kDa),^4^ cryo-electron tomography (cryo-ET) molecules and complexes that are smaller than ca. 500 kDa are often challenging to visualize, particularly in the context of densely populated cellular and tissue environments. Thus, despite advances made in the detection of ribosomes,^5^ actin filaments,^6^ microtubules,^7^ nuclear pore and basket complexes by cryo-ET,^8,9^ and the promise of template matching methods,^10^ there remains a vast number of molecules and assemblies that presently are unable to be localized by cryo-ET.

Detection, localization, and ultimate visualizations of biological molecules that are not immediately discernable by electron microscopy have benefited from the use of electron dense fiducials, beginning with the use of iron-loaded ferritin bound to antibodies.^11^ Since then, gold nanoparticles^12^ (AuNPs) conjugated to ‘target binders’, such as antibodies, antibody fragments, or small molecule ligands, have emerged as powerful tools in electron microscopy.^13^ The development of methods to synthesize homogeneous and water-soluble AuNPs, from ca. 1 nm to 3 nm in diameter,^14–18^ together with approaches to conjugate the AuNPs to single cysteine residues on the target binder^19,20^ via the Murray place exchange reaction,^21^ have further expanded the use of AuNP conjugates.

Despite these advances in the synthesis and conjugation of AuNPs, the use of AuNP conjugates to precisely localize target complexes has been hobbled by conjugation of AuNPs to the target binders via flexible linkers. For AuNPs to more accurately define the location of the target complex, we have explored methods to directly couple small and homogeneous AuNPs to defined elements of protein structure on Fab fragments that bind to 3D epitopes on a target AMPA receptor complex. By rigid and precise coupling of the AuNP to cysteine residues displayed on specific elements of secondary structure on the Fab constant domain, we can ‘program’ Fabs to harbor single AuNP binding sites, thus yielding distinct AuNP positions in the resulting AuNP –Fab target complex. Because the AuNP is rigidly fixed to the Fab, the AuNP can be exploited for target molecule localization with sub nanometer precision, for guiding particle pose in subtomogram averaging (STA), and for probing sub nanometer conformational changes in AMPA receptors at synapses.

## Results

### Anti-GluA2 15F1 Fab construct design for AuNP coupling

To develop a model system for AuNP conjugation and target protein labeling, we employed the anti GluA2 15F1 Fab^22^ and GluA2-containing AMPA receptors from mouse brain.^23,24^ The 15F1 antibody and its Fab fragment have been widely used for super resolution microscopy of AMPA receptors at synapses,^22,25^ for labeling of AMPA receptors within neurons, where it does not impact receptor function or localization,^26^ for immunoaffinity isolation of native AMPA receptors,^23^ and for use as a fiducial label for single particle cryo-EM studies.^27^ The amino acid sequence of the 15F1 antibody is known^23^ and constructs of the Fab are amenable to large-scale expression and purification. Even though we have obtained medium resolution reconstructions of the 15F1 Fab from single particle studies of AMPA receptor –Fab complexes, and the Fab structure can be predicted, we first determined an experimental, high-resolution x-ray crystal structure of the 15F1 Fab (**Fig. 1a, Supplementary Fig. 1, Supplementary Table 1**).

**Fig. 1.**
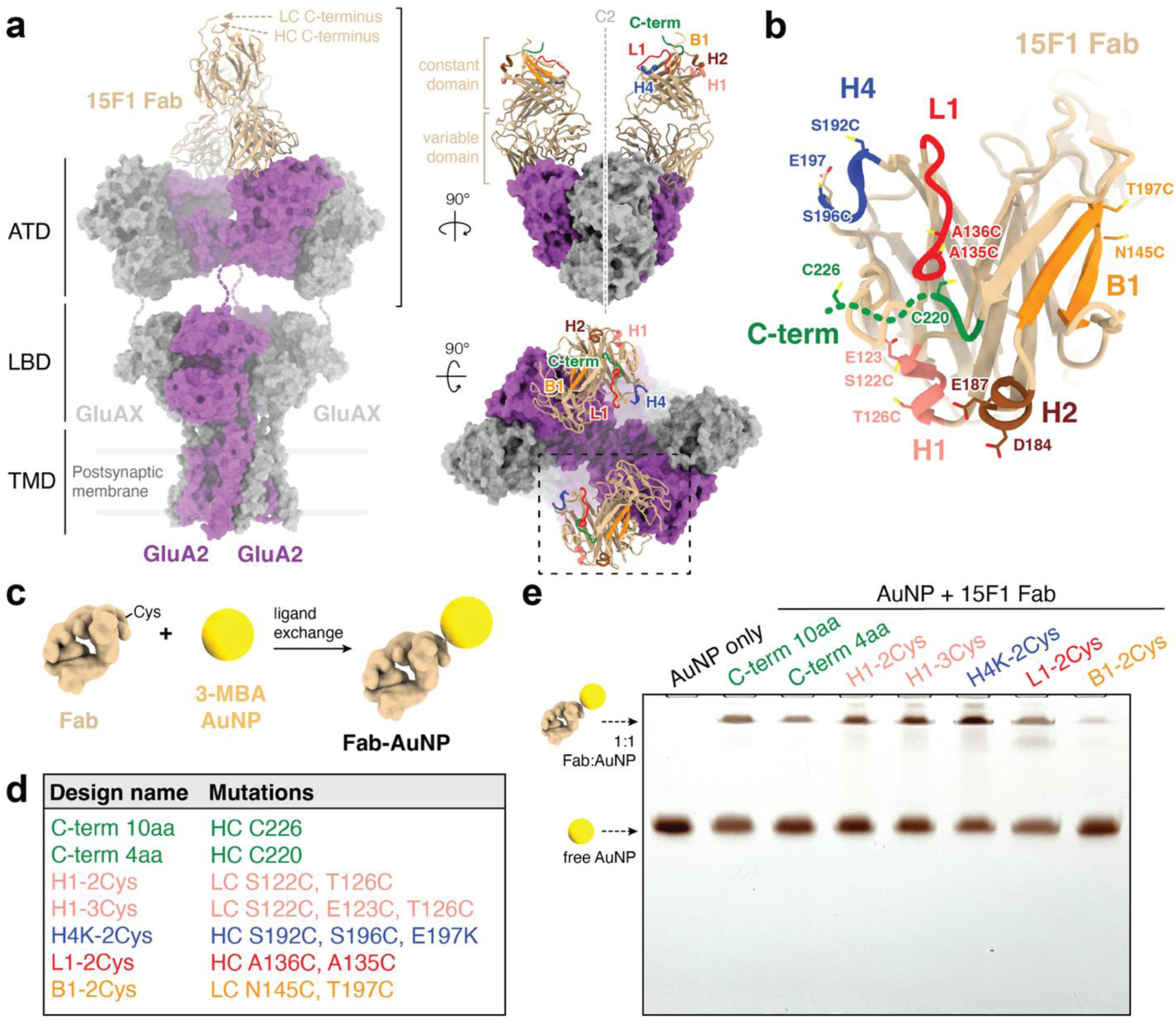
AuNP conjugation to discrete regions of the 15F1 Fab constant domain. **a**, Structure of native AMPA receptor bound to anti-GluA2 15F1 Fab (PDB ID 7LDD with aligned 15F1 Fab crystal structure PDB ID 38DC). **b**, Top-down view of the 15F1 Fab crystal structure with constant domain regions denoted and mutated cysteine residues modeled. **c**, Schematic of site-specific conjugation between reduced 15F1 Fab and 3-MBA protected AuNPs to generate 15F1 Fab-AuNP conjugates. **d**, List of cysteine 15F1 Fab mutants. HC, heavy chain; LC, light chain. **e**, Native PAGE gel comparing AuNP conjugation efficiency across a set of 15F1 cysteine mutants.

Our choice of sites on the 15F1 Fab for labeling by AuNPs was predicated on the notion that we wanted sites that would rigidly bind the AuNP without interfering with Fab binding to the receptor, by way of direct steric clashes, or by interactions between multiple AuNPs bound to a receptor complex. Thus, we began by making a parent ‘C-term 10 aa’ construct^27^ containing a typical ‘flexible’ attachment site at a cysteine located 10 residues after the last well-structured residue of the heavy chain C-terminus (**Fig. 1b-d**). Next, we simply shortened the flexible C-terminus of the heavy chain from 10 amino acids to 4, yielding the ‘C-term 4 aa’ construct, which we expected to reduce linker flexibility for monovalent C-terminal AuNP conjugation. Because we hypothesized that multivalent AuNP coordination may result in more rigid and well-defined conjugation, we inspected the x-ray diffraction and cryo-EM structures of AuNPs.^27–29^ We observed that the spacing between the thiol groups of the mercaptobenzoic acid molecules coordinating the gold cluster ranged from 4.5-6 Å. Because this sulfur –sulfur distance ‘brackets’ the distance between sulfur atoms on cysteine residues at positions i and i +4 on an α-helix,^30^ we hypothesized that a potential site for rigid AuNP conjugation could be defined by two cysteine residues separated by one turn on an α-helix. We thus chose to make cysteine mutations at light chain residues S122 and T126 on a short α-helix deemed ‘H1’ on the ‘flank’ of the constant domain, near the C-termini of the heavy and the light chains (H1-2Cys). We also mutated two residues (S192C, S196C) on a single turn of helix ‘H4’ on the constant domain, yielding the H4-2Cys construct. When the Fab is bound to an intact receptor, the H4 helix is positioned closer to the C2 axis in comparison to the H1 helix. Lastly, we picked two additional sites for AuNP coordination, one on a β-strand (B1-2Cys; N145C, T197C), where the sulfurs on the two cysteine residues are predicted to be about 6-7 Å apart, and another site on a ‘loop’, where the sulfur atoms on the cysteine residues are estimated to be about 6-8 Å from one another (L1-2Cys; A135C, A136C). With these variants in hand, we then determined their AuNP conjugation efficiency (**Supplementary Fig. 2-4, Supplementary Table 2**).

AuNP conjugation was carried out by way of a thiol exchange reaction, where reduced cysteine residues on the Fab replace surface-bound 3-MBA ligands on the AuNP to form stable Au-S bonds and site-specific Fab-AuNP conjugates (**Fig. 1c**). Reactions of the C-terminal constructs (10 aa and 4 aa linkers) show that shorter linker length reduces conjugation efficiency, likely due to decreased cysteine exposure and limited access to the AuNP surface (**Fig. 1e**). The H1 and H4 cysteine designs show the strongest and sharpest conjugate gel bands, which supports our hypothesis that i to i+4 spacing on α-helices enables bidentate coordination and robust conjugation efficiency. Interestingly, the trivalent thiol H1-3Cys Fab shows slightly enhanced conjugation in comparison to the divalent H1-2Cys Fab. The loop (L1-2Cys) species shows reduced conjugation, perhaps because the underlying main chain is more conformationally mobile in comparison to H1 and H4, thus disfavoring thiol engagement with the AuNP. The β-sheet (B1-2Cys) construct exhibits a poor conjugation yield, which may be due to the thiol groups forming a stable and rapidly reforming disulfide, thus precluding efficient AuNP conjugation. Alternatively, there may be other factors, such as unfavorable steric interactions surrounding the site, or the exposure of the thiol groups themselves, that disfavors AuNP conjugation.

### Local electrostatic environment influences AuNP conjugation

We speculated that thiol-mediated conjugation of AuNPs to the Fab constructs could be improved by creating a positive local electrostatic environment around the cysteine residue(s) in order to promote favorable association of the negatively charged AuNPs, akin to prior studies on scFv –AuNP conjugates.^19^ For the H1 helix, the H1-2Cys design (C122, C126) contains a nearby acidic residue (E123), which may introduce electrostatic repulsion with the negatively charged 3-MBA coated AuNP surface, reducing conjugation efficiency. To address this, E123 was mutated to alanine, cysteine, or lysine. Introducing a positively charged residue significantly improved conjugation, with the K123 mutation showing highest conjugation efficiency (**Fig. 2a**). The A123 substitution moderately increased conjugation efficiency, whereas the C123 substitution, which may participate in tridentate coordination or simply reduce the negative electrostatic potential, modestly improves conjugation efficiency relative to the A123 mutant. We further examined nearby negatively charged residues around the H1 helix, noting that helix 2 (H2) has two negatively charged residues, D184 and E187. These residues were mutated to positively charged lysine residues to increase local positive charge; however, the mutants did not produce substantial improvements in conjugation efficiency, indicating that electrostatic modulation farther from the H1 helix has a limited effect. Similar charge modulation at the H4 helix, mutating E197 to K, enhances conjugation at the H4-2Cys site, reinforcing the conclusion that local electrostatic tuning can improve AuNP conjugation (**Fig. 2b**).

**Fig. 2.**
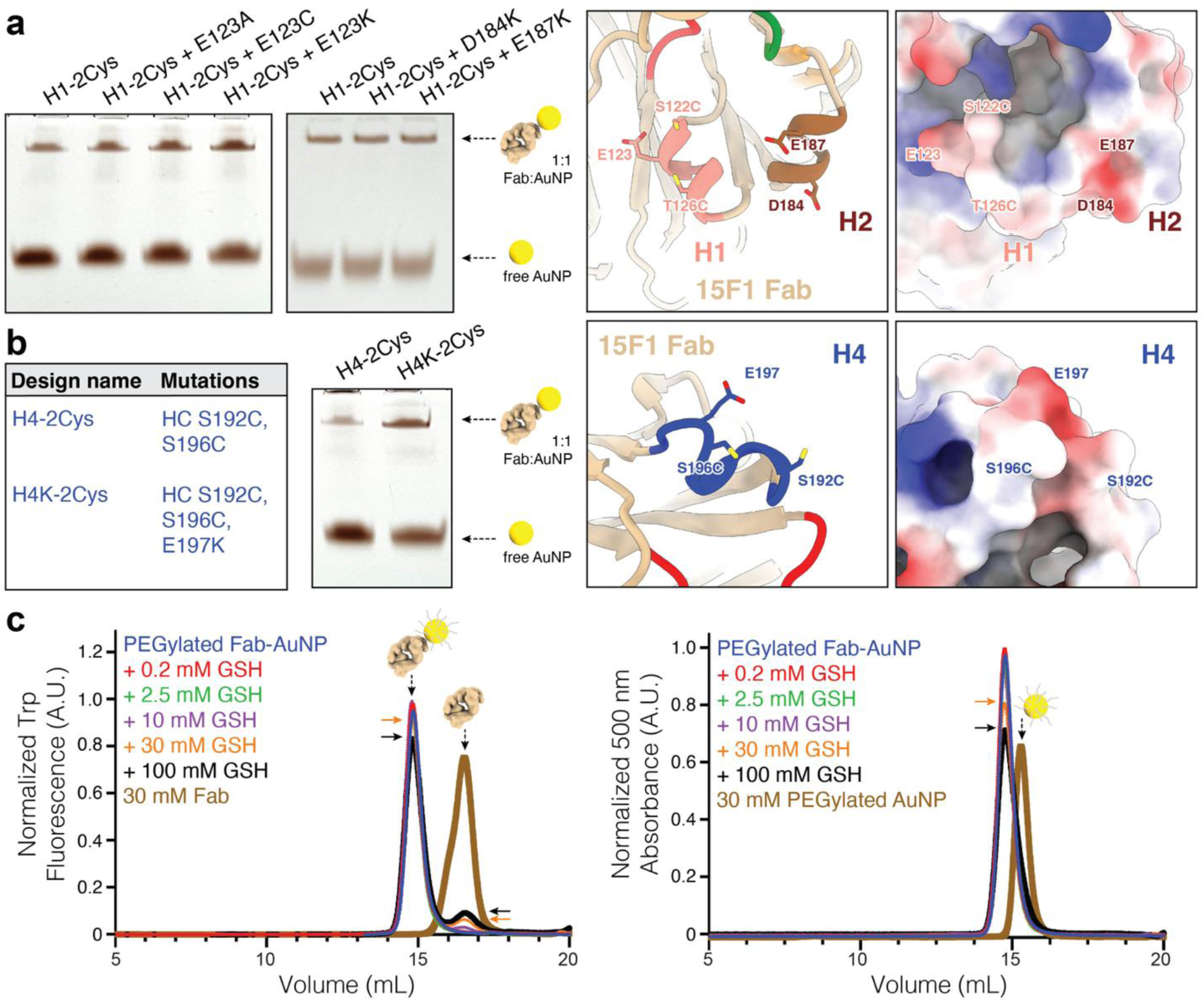
Local charge modulates reactivity of glutathione-stable conjugates. **a**, Local electrostatic effects on thiol-AuNP conjugation were assessed by mutating residue E123 (adjacent to H1-2Cys site) to alanine, cysteine, or lysine (left panel). Additional charge optimization was tested by mutating local residues D184K and E187K (right panel). **b**, Similar electrostatic optimization at the H4 site by introducing E197K mutation in the H4-2Cys background (resulting in H4K-2Cys). **c**, Stability of 15F1 C-term 10 aa Fab-AuNP conjugates under physiological reducing conditions was assessed by FSEC Trp fluorescence and AuNP absorbance at 500 nm.

### Fab-AuNP complexes are stable under physiological reducing conditions

The stability of Fab-AuNP conjugates is essential for their use in thiol-rich biological environments where Au–S interactions can be challenged by endogenous reducing agents, such as glutathione. To confer favorable behavior in solution and in tissue, we PEGylate AuNPs with saturating concentrations of PEG550-SH after Fab conjugation. We thus set out to examine the stability of these PEGylated Fab-AuNP conjugates under physiologically relevant reducing conditions using reduced L-glutathione. Fluorescence-detection size-exclusion chromatography (FSEC)^28,29^ analysis using both Trp fluorescence and AuNP-specific 500 nm absorbance channels shows that the Fab-AuNP C-term 10 aa conjugate, which contains a single cysteine conjugation site, retains a monodisperse peak in presence of 0.2–100 mM GSH (**Fig. 2c**). Minimal dissociation was observed at 10 mM GSH, consistent with stability close to physiological conditions. At higher GSH levels, a slight increase in free Fab is observed, suggesting partial destabilization. A similar analysis was performed with the H4K-2Cys conjugate, which showed a comparable monodisperse profile across the same GSH range, consistent with conjugate stability under reducing conditions (**Supplementary Fig. 5**). Overall, the majority of Fab-AuNP conjugate remains intact, supporting the suitability of the conjugates for labeling in cellular environments (**Fig. 2c**).

### Structural characterization of AMPA receptor –Fab-AuNP complexes

To assess how the coupling of AuNPs to different regions of the Fab constant domain affects the order and localization of the AuNP relative to the constant domain, how it defines the positions of multiple AuNPs within the receptor complex, and the extent to which the AuNPs impact resolution of the receptor portion of the complex, we determined structures of the 15F1 Fab-AuNP conjugates (C-term 10 aa, C-term 4 aa, H1-2Cys, H1-3Cys, L1-2Cys, and H4K-2Cys) bound to native AMPA receptors by single particle cryo-EM (**Supplementary Table 3**). To achieve this, we immunoaffinity purified endogenous AMPA receptor assemblies from mouse brain tissue in the presence of a saturating concentration of anti-GluA2 15F1 Fab-AuNP (C-term 10 aa and C-term 4 aa) conjugate or with the anti-GluA1 11B8scFv and 15F1 Fab-AuNP (H1-2Cys, H1-3Cys, L1-2cys, and H4K-2Cys) conjugates (**Fig. 3a, Supplementary Fig. 6**). During both 2D and 3D classification, the high contrast AuNPs drove particle alignment, yet clear receptor density was visible adjacent to the AuNPs in many classes (**Supplementary Fig. 7-8**). Variants with multivalent conjugation sites on elements of secondary structure, such as on the H1 or H4 helices, resulted in more well-defined receptor features in both 2D and 3D classes, consistent with more rigid AuNP attachment in these conjugates (**Fig. 3a**). In agreement with previous native AMPA receptor structural studies, the 2:1 15F1 Fab-AuNP:AMPA receptor species with GluA2 in the B and D positions was the most abundant class for all but one variant.^23,24^ In the L1-2Cys dataset, only a 1:1 15F1 Fab-AuNP:AMPA receptor species was observed, suggesting that the conjugation of AuNPs to the L1 loop results in steric clashes that are incompatible with a 2:1 bound Fab-AuNP:AMPA receptor assembly. Indeed, in the L1-2Cys Fab-bound receptor, we estimate the distance between the centroids of the conjugation cysteine Cα atoms to be 6.1 nm. Thus, we conclude that an inter-AuNP conjugation site spacing of less than 6 nm, in the context of the sizes of AuNP and PEG used for our conjugate preparations, is incompatible with saturated labeling.

**Fig. 3.**
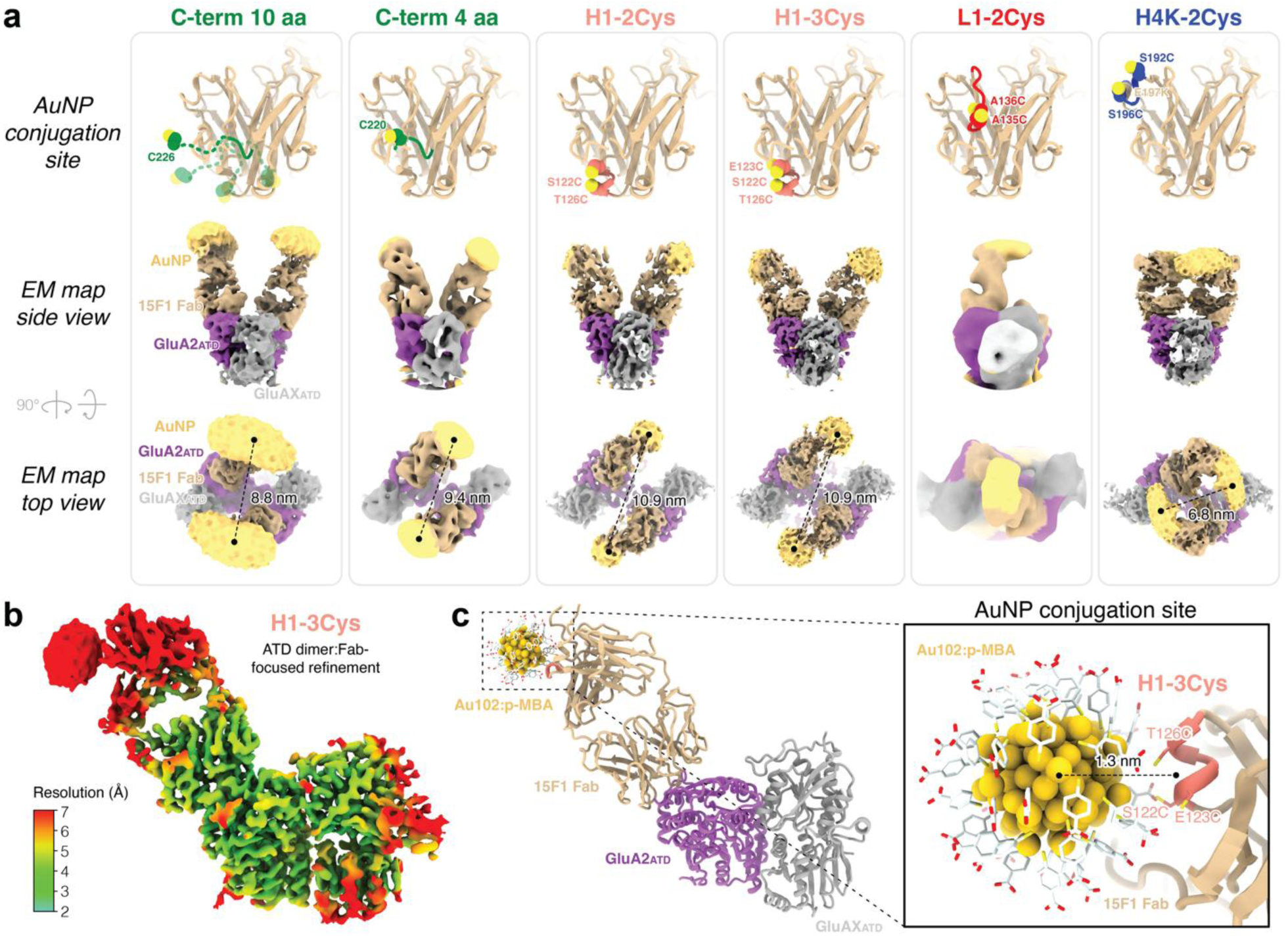
Rigidities of AuNP-Fab conjugates vary across conjugation sites. **a**, AuNP conjugation site cysteines in 15F1 Fab constant domain (top panel). Side (middle panel) and top (bottom panel) views of the single particle cryo-EM map (ATD:Fab-focused) of each 15F1 Fab-AuNP:native AMPA receptor complex. **b**, Local resolution map (ATDdimer:Fab-focused) for H1-3Cys 15F1 Fab-AuNP complex. **c**, Docked model of H1-3Cys ATDdimer:15F1 Fab-AuNP:3-MBA into map shown in (b), with the distance between the center of the AuNP density and the conjugation site denoted.

The structures of 2:1 immunolabeled receptor complexes were determined for each variant by refining the full complex, signal subtracting the AuNP densities, refining the subtracted particles, and then reconstructing the complex without alignment using the un-subtracted, AuNP-containing particle set. This resulted in Fab-AuNP bound receptor reconstructions at 7-12 Å resolution (**Supplementary Table 3**). Local focused refinement of the C2 symmetric Fab-amino terminal domain (ATD) region resulted in improved 4-8 Å resolution reconstructions with clearly defined AuNP density adjacent to the installed conjugation cysteines for each variant (**Fig. 3a**). The C-term 10 aa variant displayed the most diffuse AuNP density ‘cloud’, consistent with the expected flexibility of the AuNP conjugation site at the end of a 10-residue unstructured linker. This flexibility was significantly reduced by removing 6 residues from the linker in the C-term 4 aa variant, which resulted in a smaller AuNP density feature positioned closer to the Fab. The L1-2Cys and H4K-2Cys variant AuNP densities were slightly oblong but less than half the size of the AuNP density from C-term 10 aa, indicating that the L1 loop and H4 helix are only slightly mobile structural elements. In contrast, the AuNP density for either the H1-2Cys or H1-3Cys variants were nearly spherical and smaller relative to the other variants, indicating rigid coupling of the AuNP.

Symmetry expansion and local refinement of a single ATD_dimer_:Fab further improved local reconstructions to 3.9-7.0 Å. For the largest dataset, H1-3Cys, the local ATD_dimer_:Fab-AuNP region was resolved to 3.5 Å after local CTF refinement and motion correction, allowing for confident docking of GluA2 ATD, 15F1 Fab, and AuNP models into the density (**Fig. 3b-c**). A previously determined crystal structure of a para-MBA coated AuNP fit well to the overall shape and size of the AuNP EM density.^30^ Limited resolution in this region of the map precluded visualization of the cysteine side-chain density at the AuNP conjugation site, potentially due to artifacts from the AuNP signal and intrinsic flexibility of the Fab Fc region. However, based on the 15F1 Fab crystal structure docking, the AuNP density center is approximately 1.3 nm from the α-carbon centroid position of the three conjugation cysteines on helix H1, consistent with site-specific, rigid coupling to the Fab (**Fig. 3c**). Comparison of the AuNP density in the focused ATD_dimer_:Fab-AuNP maps for the H1-2Cys and H1-3Cys variants reveals a 0.2 nm shift in the AuNP density center in the direction of the added E123C mutation, consistent with the 0.2 nm shift in mean position of the H1-2Cys (S122C and T126C) or H1-3Cys (S122C, E123C, and T126C) conjugation residue centroids (**Supplementary Fig. 9**). Although the reconstruction resolution precludes definitive assignment of which cysteines are actively conjugated to the AuNP, this observation suggests that the third cysteine in H1-3Cys may participate in conjugation, or that removal of the negatively charged glutamic acid residue results in a slight shift of the conjugated AuNP position towards the C123 residue.

### Fab-AuNP conjugates enable subtomogram averaging

To test whether the Fab-AuNP conjugates are compatible with STA, we collected 186 tilt series on the purified H4K-2Cys 15F1 Fab-AuNP:native AMPA receptor complex. Although the H4K-2Cys conjugate displayed moderately rigid coupling by single particle analysis, we expected that reducing the AuNP pair spacing in the receptor complex may help differentiate adjacent receptors *in situ*. We identified AuNP positions within the tomograms and applied a distance-based cutoff to select isolated pairs of AuNPs within 5-9 nm, indicative of native AMPA receptors with two GluA2 subunits in the B/D positions (**Fig. 4a, Supplementary Table 4**). Indeed, upon manual inspection of particles picked using this strategy, clear receptor density was observed next to each of these AuNP pairs. Using this approach, 5,975 putative particle positions were picked and analyzed using the Warp pre-processing^31^ and RELION 5^32^ STA pipelines. An initial round of refinement produced an AuNP-focused map with reasonably well-resolved receptor density (**Supplementary Fig. 10a**). After recentering on the receptor density and separate focused refinements on the ATD and ligand-binding domain:transmembrane domain (LBD:TMD) regions, we attained 15-20 Å reconstructions of an H4K-2Cys 15F1 Fab-AuNP labeled native AMPA receptor (**Fig. 4b-f, Supplementary Fig. 10a-d**). These maps are consistent with our single particle reconstruction of the same complex but were notably processed without the need for particle classification, clean-up, or AuNP subtraction. To test whether the AuNPs limit target alignment despite local masking, we erased the AuNP signal from the raw tilt images, deriving the positions from the AuNP positions within the tomogram, and ran identical masked refinements on the AuNP-erased particle set. This resulted in a reconstruction with only marginally improved receptor density (**Supplementary Fig. 10e-f**). Based on these STA analyses, we expect that the rigidly coupled AuNPs are compatible with STA approaches and can drive initial alignment of small macromolecular complexes. Subsequent bootstrapping from these initial AuNP-driven alignments with AuNP-subtracted data may improve reconstruction quality, particularly for regions of a complex farther from the AuNP or that are separated by flexible regions.

**Fig. 4.**
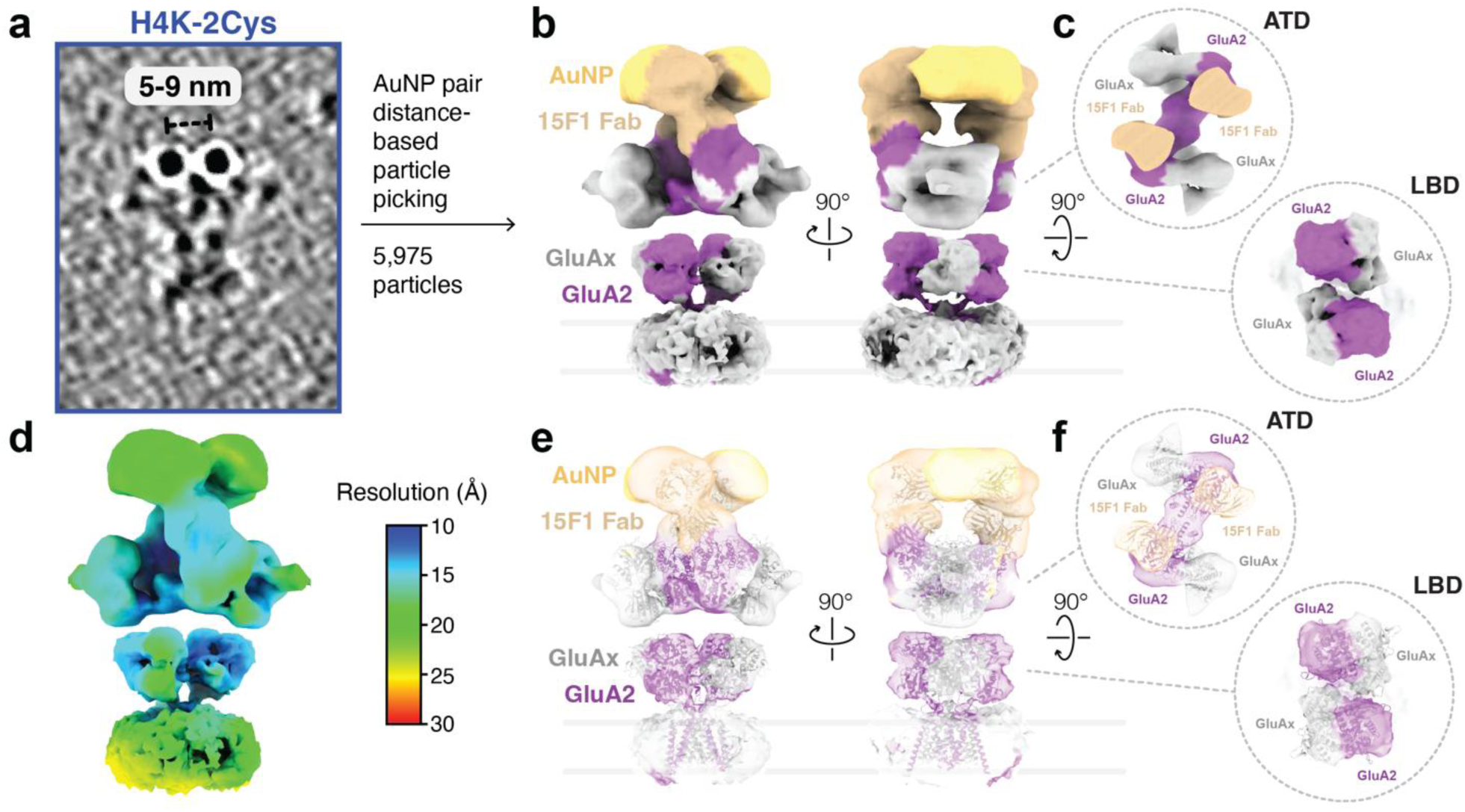
AuNP-guided particle picking streamlines STA workflow. **a**, Example H4K-2Cys 15F1Fab-AuNP:native AMPAR particle in H4K-2Cys tilt series dataset. **b**, Composite STA map of H4K-2Cys 15F1Fab-AuNP:native AMPAR complex from 5,975 particles. **c**, Clipped views of ATD and LBD receptor regions in map from panel **b**. **d**, Local resolution STA map of H4K-2Cys 15F1-AuNP:native AMPAR complex. **e-f**, Overlays of STA map and clipped views from panels **b-c** with 15F1 Fab:native AMPA receptor structure (PDB ID 7LDD).

### Fab-AuNP label defines AMPA receptor position and orientation at synapses

One goal for use of the rigid 15F1 Fab-AuNP conjugates is unambiguous identification of the position and orientation of individual AMPA receptors at native glutamatergic synapses. This is particularly difficult within the crowded synaptic environment, and is made even more challenging in native brain tissue, which requires cryoprotectants for efficient freezing, resulting in lower contrast than an ideal sample. Additionally, AMPA receptors are known to cluster at native synapses,^33–35^ further complicating AuNP-receptor mapping with closely spaced receptors. To assess whether receptors could be precisely localized *in situ* with the rigid 15F1 Fab-AuNP conjugates, we employed our established brain tissue-derived synapse cryo-ET imaging pipeline^27^ and tested receptor labeling with three of the variants: C-term 10 aa, H1-2Cys, and H4K-2Cys.

For each variant, nearest neighbor distances between synaptic AuNPs were compared with the intra-receptor AuNP pair distances from our single particle reconstructions (**Fig. 5a-b**). For the C-term 10 aa and H1-2Cys variants, the mean and standard deviation of the Gaussian fits of the nearest neighbor distributions were 7.4 ± 1.8 and 8.1 ± 2.4 nm, respectively. These averages are lower than the center-to-center distance between the AuNP densities observed in the respective single particle EM maps, likely because the nearest neighbor is often from a closely positioned adjacent receptor, rather than from the intra-receptor AuNP pair. By contrast, the nearest neighbor distribution of the H4K-2Cys variant was less broad, with a Gaussian fit of 7.0 ± 1.2 nm. This is in close agreement with the expected intra-receptor pair distance of 6.8 nm observed in the H4K-2Cys single particle and STA maps and suggests that in most cases, the nearest neighbor of each AuNP is the intra-receptor AuNP pair.

**Fig. 5.**
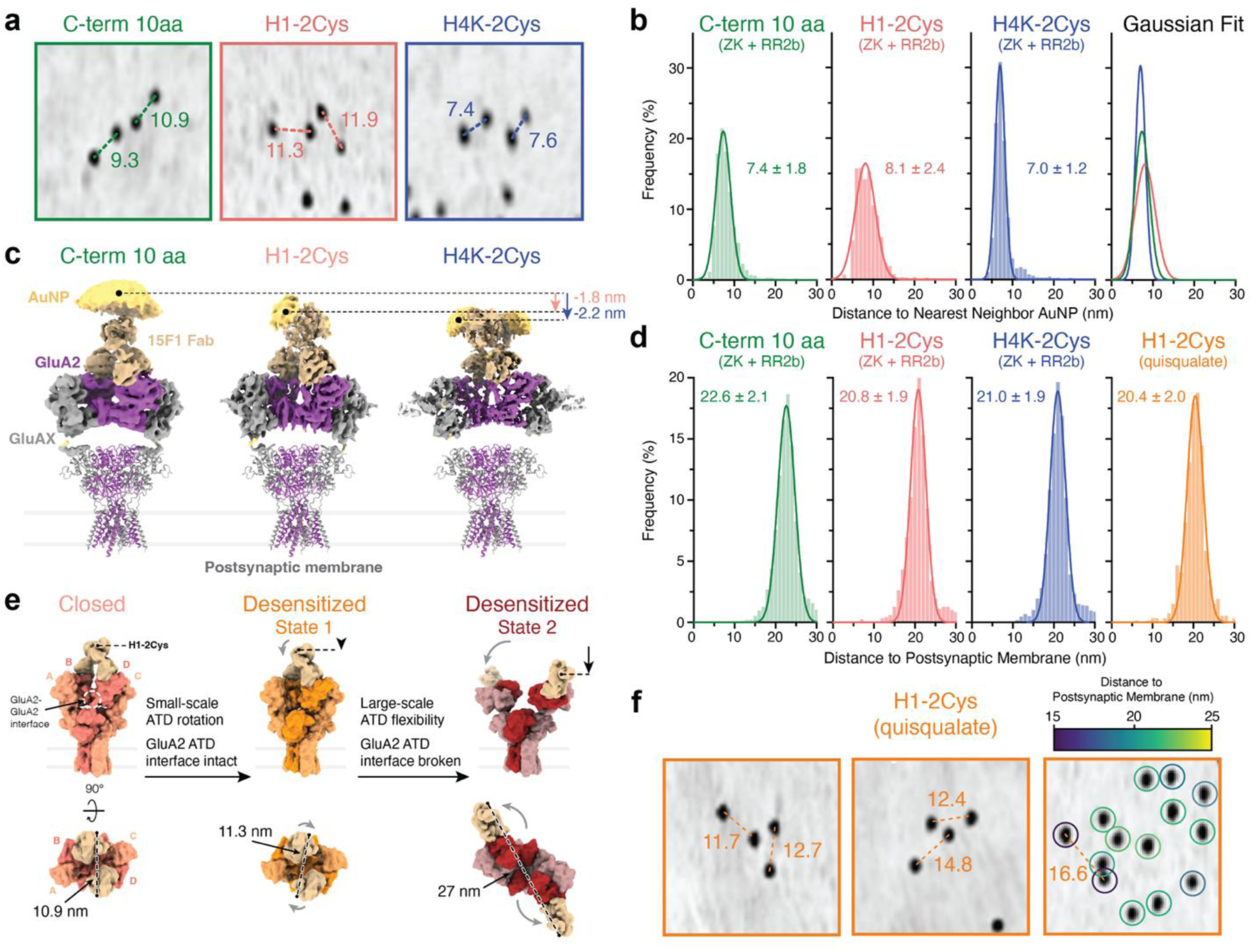
AuNP label positions and *in situ* receptor conformations. **a**, Minimum intensity projection views of small AuNP clusters at synaptic cleft, with distance between putative intra-receptor AuNP pairs denoted. **b**, Nearest neighbor distances of AuNPs at synapses *in situ* for each conjugate, with Gaussian fit mean and standard deviation shown (C-term 10 aa n=5,566; H1-2Cys n=1,184; H4K-2Cys n=3,701). **c**, Differences in predicted AuNP to postsynaptic membrane distances for three AuNP conjugates based on ATD:15F1 Fab-focused single particle cryo-EM maps. **d**, AuNP to postsynaptic membrane distances for AuNPs at synapses in situ, with Gaussian fit mean and standard deviation shown (C-term 10 aa n=5566; H1-2Cys n=1184; H4K-2Cys n=3701; H1-2Cys + quisqualate n=512). **e**, Models of AMPA receptors in a closed state (PDB 7LDD) and in two previously observed ATD-splayed states (based on PDB 7RZ7 and EMDB 44245) that may occur during receptor desensitization. **f**, Minimum intensity projection views of synaptic cleft regions from H1-2Cys labeled quisqualate sample with putative intra-receptor AuNP pair and postsynaptic membrane distances denoted.

For each of the labels, we also looked at minimum intensity projections of synaptic cleft regions and examined small clusters of AuNPs containing pairs that could be unambiguously assigned to individual receptors (**Fig. 5a, Supplementary Fig. 11**) under the assumption that most receptors contain two GluA2 subunits in the B and D positions.^23^ In most of these cases, estimated AuNP pair distances are within 2 nm of that predicted by single particle analyses. In closely packed AuNP clusters, the nearest neighbor AuNP for the H4K-2Cys variant is much more frequently a clearly identifiable intra-receptor pair than in either the C-term 10 aa or H1-2Cys variants, in agreement with the nearest neighbor distance analysis. Therefore, for AMPA receptor labeling at native synapses, the H4K-2Cys variant most accurately defines individual receptor positions with nearest neighbor pairs, particularly in large and densely packed receptor clusters.

### AuNP positions directly define AMPAR conformational states *in situ*

As a prelude to determining whether AuNP positions could be used as a probe of receptor conformation, we next examined the AuNP to postsynaptic membrane distance for each variant in cryo-electron tomograms of Fab-AuNP labeled brain-derived synapses. Based on our single particle cryo-EM maps, the center of the AuNP densities for the H1-2Cys and H4K-2Cys variants are 1.8 and 2.2 nm closer to the receptor transmembrane domain than the C-term 10 aa variant (**Fig. 5c**). At synaptic clefts, the Gaussian fit mean of AuNP position to postsynaptic membrane distance for the H1-2Cys (20.8 ± 1.9 nm) and H4K-2Cys (21.0 ± 1.9 nm) variants were 1.8 and 1.6 nm lower, respectively, than for the C-term 10 aa label (22.6 ± 2.1 nm) (**Fig. 5d**). These results from the cryo-ET experiments are consistent with the AuNP positions observed in our single particle analyses and confirm that single digit nanometer differences in average AuNP positions can be resolved *in situ*.

Armed with the knowledge that our most rigidly coupled 15F1 Fab-AuNP conjugate, H1-2Cys, forms a well-defined assembly with native GluA2-containing AMPA receptors, we set out to probe *in situ* conformational changes of the receptor at specific steps in the gating cycle. Early single particle analysis of native AMPA receptors suggested large-scale conformational changes upon receptor desensitization.^36^ More recently, x-ray crystal structures and additional single particle studies show that AMPA receptors undergo a spectrum of conformational motions upon desensitization, from relatively small-scale rotation of the ATD and displacement of the ATD toward the receptor transmembrane domain, to larger-scale rearrangements accompanied by splaying of the ATD layer and rupture of the ATD B-D interface.^37–41^ Because these conformational changes result in a relative movement of the ATD closer to the receptor TMD, and the 15F1 Fab binds to the ATD, we sought to measure the change in distance between the AuNPs bound to the 15F1 Fab and the postsynaptic membrane, in antagonist-bound/resting and agonist-bound/desensitized states (**Fig. 5e**).

To stabilize receptors in a closed, non-desensitized state, we included the potent competitive antagonist ZK200775 (ZK)^42^ and the positive allosteric modulator (R,R)-2b^43^ during brain slice preparation for each of the C-term, H1-2Cys, and H4K-2Cys variants. In a separate preparation with the H1-2Cys variant, we included 1 mM quisqualate, a high-affinity agonist that stabilizes receptors in a closed, desensitized state.^44^ The Gaussian fit mean of AuNP position to postsynaptic membrane distance for the desensitized-state H1-2Cys labeled receptors with quisqualate (20.4 ± 2.0 nm) was 0.4 nm smaller than the closed-state H1-2Cys labeled receptors with ZK and RR2b (**Fig. 5d**). AuNP pair distances for putative receptor-bound pairs in quisqualate samples were similar or only slightly larger (∼1-3 nm) than those observed in ZK + (R,R)-2b samples (**Fig. 5f**). This suggests that in most receptors, desensitization is associated with relatively small ATD rearrangements and movements towards the postsynaptic membrane. However, we note that a few of the AuNPs are positioned at least 4 nm closer to the postsynaptic membrane than the average in both the ZK + (R,R)-2b and quisqualate datasets. These could represent a subset of synaptic receptors with GluA2 subunits in an A/C position, receptors with substantially splayed ATDs, or incompletely assembled monomeric or dimeric receptor assemblies.

## Discussion

Here we show that AuNPs can be coupled to α-helical elements of secondary structure on the constant domain of a Fab fragment, utilizing cysteine residues at positions i and i+4, yielding rigidly and robustly bound AuNPs. The structural elements we surveyed for conjugation are conserved across antibody constant domains from common host species used for antibody development (**Supplementary Fig. 12**), making the conjugation sites defined in this study well-suited for application to equivalent sites on other Fab fragments.^45^ We observe that the conjugation efficiency of 3-MBA protected AuNPs, which have a negatively charged surface, is influenced by the electrostatic environment around the cysteine residues, with nearby negative charge reducing and positive charge improving conjugation efficiency, respectively. This effect is observed only in the immediate vicinity of the conjugation cysteines, as introduction of positively charged residues on an adjacent α-helix, 1 nm away from the conjugation site, had no readily discernable effect on AuNP conjugation. Therefore, neutralization or charge swap of negatively charged residues near intended conjugation cysteines should be considered when engineering AuNP coupling sites.

To assess the relative rigidity of AuNPs within these Fab fragment conjugates and the positioning of AuNPs within target-bound complexes, we determined the structures of a representative panel of Fab-AuNP conjugates in complex with purified native AMPA receptors. These receptor-bound conjugates displayed varying extents of AuNP flexibility, with the most rigid conjugates resulting from AuNPs coupled to at least two cysteines on helix ‘H1’. Notably, the more rigidly coupled AuNP conjugates reduced the necessity for AuNP signal subtraction to attain well-aligned target reconstructions by single particle cryo-EM. These analyses demonstrate that target regions directly bound to Fab-AuNP labels can be resolved to 3-4 Å resolution, despite their proximity to the high contrast AuNPs, and provide the highest resolution structures of AuNP-protein conjugates currently available.

Although antibody, antibody fragment, and small molecule ligand AuNP conjugates,^46^ as well as genetically encoded multimeric tags (GEMs)^47^ and DNA signpost origami tags (SPOTs),^48^ have been successfully employed for cryo-ET labeling and target localization in previous work, flexible linkers between the identifiable label feature and molecular target have limited their utility. In most cases, identification of target position and orientation is not directly interpretable from these label positions without further processing steps such as STA. Additionally, STA of previous AuNP-labeled targets has required suppression of the AuNP signal to attain target-focused particle alignments.^46^ Using one of our Fab-AuNP labels, we show that an AuNP-guided particle picking approach robustly identifies targets in a purified Fab-AuNP labeled “test” AMPA receptor sample, without the need for any additional particle curation or classification. From a set of approximately 6,000 particles, AuNP-labeled receptors can be reconstructed to 15-20 Å by STA, without erasing the AuNP signal. As a result, these rigid AuNP conjugates can function both as target identifiers and as fiducials that drive initial alignment of small macromolecular targets for STA.

By pairing our previously developed high-pressure frozen brain tissue lamella preparation and synapse-targeting workflow^27^ with our rigidly coupled Fab-AuNP conjugates, we show that we can label AMPA receptors at synapses *in situ*, map the AuNP positions within tomograms, and identify AuNP pairs bound to individual receptors at synapses. For AMPA receptors, we find that the coupling site resulting in the closest intra-receptor AuNP pair spacing, H4K-2Cys, resolves individual receptors most clearly within the tightly packed receptor clusters observed at synapses *in situ*. This site-specific rigid AuNP coupling also facilitates direct exploration of target structural changes based solely on target-bound AuNP positions, which can be interpreted as movements of epitope-containing domains relative to other tomogram landmarks, such as membranes, without the need for STA approaches that typically require thousands of particles.^49^ Using this approach, we measured AuNP-to-postsynaptic membrane distances and observed conjugation site-specific single digit nanometer changes in relative AuNP-membrane distances. Using the most rigidly coupled H1-2Cys conjugate, we show that a small movement of the receptor ATD towards the postsynaptic membrane accompanies receptor desensitization *in situ* for most GluA2-containing receptors. We expect that our atomically precise AuNP coupling approach will expand the accessibility of cryo-ET technologies for smaller and historically inaccessible molecules and complexes *in situ*.

## Methods

### Synthesis of monomeric AuNPs

Gold (III) chloride trihydrate (Oakwood Chemical, Cat. No. 386720), 3-mercaptobenzoic acid (Sigma-Aldrich, Cat. No. 451436), sodium hydroxide (Fisher Scientific, Cat. No. BP359212), sodium borohydride (Sigma-Aldrich, Cat. No. 480886), sodium chloride (Fisher Scientific, Cat. No. S271), and methanol (Fisher Scientific, Cat. No. BP11054) were used. Deionized water (Milli-Q grade) was used for all aqueous solutions.

AuNPs utilized for preparation of the anti-GluA2 15F1 Fab-AuNP conjugate containing a free cysteine at heavy chain C-terminal residue 226 (‘C-term 10 aa’) were synthesized as previously described,^27^ using a 7:1 molar ratio of 3-mercaptobenzoic acid (3-MBA) to gold (III) chloride trihydrate (HAuCl_4_·3H_2_O), producing AuNPs of approximately 2.2 nm diameter. Briefly, in a 15 mL plastic (Falcon) tube, 1 mL of 84 mM 3-MBA (12.75 mg/mL in methanol) and 0.4 mL of 28 mM HAuCl_4_·3H_2_O (11.03 mg/mL in methanol) were added to 3.5 mL of water, producing a white precipitate. The solution was brought to a pH of 13 by drop-wise addition of 49.5 µL of 10 M NaOH, inverting the tube to mix and resolubilize the precipitate, then rotated on a HulaMixer Sample Mixer (Thermo Fisher) at 10 rpm for at least 16 hours at room temperature. The reaction was moved to a 50 mL plastic (Falcon) tube and diluted with 28.7 mL of a 27% aqueous solution of methanol. A fresh stock of 150 mM (5.67 mg/mL) sodium borohydride (NaBH_4_) was prepared in 10 mM NaOH, and 450 µL of the stock was added to the reaction, to a final concentration of 2 mM. The reaction was continued by rotation at room temperature for an additional 4.5 hours, during which the solution slowly turns light brown. AuNPs were precipitated by addition of NaCl to 100 mM, adding 694 µL of a 5 M NaCl stock, then diluted with 80 mL of methanol. The reaction was divided between three 50 mL plastic (Falcon) tubes, spun at 4,000 *g* for 20 minutes, and the resulting AuNP pellets were each gently resuspended in 10 mL of a 75% aqueous solution of methanol using a serological pipette. The three resuspended pellets from a single reaction were combined into a single 50 mL plastic (Falcon) tube and spun again at 4,000 *g* for 20 minutes. Supernatant was discarded and the pellets were dried overnight in a glass vacuum desiccator.

AuNPs used in preparation of all other conjugates were synthesized as previously described,^18^ using a 5:1 molar ratio of 3-MBA:HAuCl_4_ and producing AuNPs of approximately 1.4 nm diameter. Briefly, in a 400 mL glass beaker, 9.07 mL of 33 mM HAuCl₄·3H₂O (12.99 mg/mL in methanol) and 18.14 mL of 84 mM 3-MBA (12.94 mg/mL in methanol) were combined and gently swirled to mix. Subsequently, 68 mL of water was added, resulting in formation of a white precipitate. The pH was adjusted to 13 by dropwise addition of 5.69 mL of freshly prepared 2 M NaOH while gently swirling the solution to fully resolubilize the precipitate. The reaction mixture was covered with aluminum foil and stirred at room temperature for 20 hours, yielding a clear, colorless solution. Following incubation, the reaction mixture was diluted with 23.21 mL of methanol and 68.28 mL of water to adjust solvent composition prior to reduction. Reduction was initiated by addition of 2.37 mL of freshly prepared 0.19 M sodium borohydride (7.18 mg/mL) in water under continuous stirring. The reaction was maintained at room temperature for an additional 5.5 hours under light-protected conditions, during which gradual development of a dark brown coloration was observed, consistent with the reduction of Au(III) species and formation of 3-MBA stabilized AuNPs. AuNPs were precipitated by increasing the ionic strength of the reaction mixture through addition of 16 mL of 0.1 M NaCl, followed by dilution with 32 mL of methanol to decrease solvent polarity and promote nanoparticle precipitation. The reaction mixture was distributed into four 50 mL conical tubes and centrifuged at 4,000 × g for 20 minutes at 4 °C to pellet the AuNPs. Supernatants were carefully removed, and the resulting pellets were briefly air-dried to eliminate residual solvents.

Dried AuNP pellets were resuspended in water to a final concentration of 5-10 mg/mL, yielding a homogeneous dark brown solution. Aliquots were transferred to light-protected (amber or foil-wrapped) microcentrifuge tubes and stored at 4 °C. Under these conditions, the AuNPs remained stable for at least one year. Each AuNP preparation was analyzed by native polyacrylamide gel electrophoresis (PAGE) to confirm size and homogeneity. Native gels were prepared using 12% (w/v) Acrylamide/Bis-acrylamide and 10% (v/v) glycerol in 0.5x TBE buffer (50 mM Tris base, 50 mM boric acid, and 1.25 mM EDTA, pH 8.3). The gel solution was degassed prior to polymerization, which was initiated by addition of 0.001% (v/v) TEMED and 0.05% (w/v) freshly prepared ammonium persulfate. Electrophoresis was performed under non-denaturing conditions to assess AuNP monodispersity, batch-to-batch consistency, and separation of Fab-AuNP conjugate species. Gels were run at 180 V for 90 minutes under cold conditions to preserve conjugate integrity and minimize diffusion.

### Expression and purification of cysteine engineered 15F1 Fab constructs

The wild-type 15F1 Fab (C-term 10 aa), retaining a 10 amino acid unstructured C-terminal sequence, was generated by cloning the light chain (LC) and heavy chain (HC) genes into the pFastBac Dual vector for expression in Sf9 insect cells. Both chains were fused to N-terminal GP64 signal peptides to enable secretion. The HC was fused with HRV 3C protease site followed by Strep-tag II for affinity purification whereas the light chain was unmodified at the C terminus. The HC C terminus contains a surface exposed reactive cysteine (Cys226), positioned upstream of an HRV 3C protease site and Strep-tag II. The cysteine variants (**Supplementary Table 2**) were generated for this study by site-directed mutagenesis using the QuikChange method (Agilent, Cat. No. 210518) (Ref: QuikChange II Site-Directed Mutagenesis Kit Instruction Manual) or by Gibson assembly (NEB, Cat. No. E2611L)^50^. In all constructs other than the 15F1 C-term 10 aa and 4 aa, the LC Cys220 was mutated to an Ala.

Recombinant P1 and P2 baculovirus for each Fab construct was generated using established baculovirus expression protocols, and viral titers were determined using the Sf9 Easy Titer cell line, as previously described.^51^ Sf9 cells were cultured at density of 2-2.5 x 10⁶ cells/mL in SF900 III SFM medium (Thermo Fisher Scientific, Cat. No. 1258027) in flat-bottom Erlenmeyer flasks prior to infection. Large scale expression was performed by infecting 3 L cultures of Sf9 cells with P2 baculovirus at a multiplicity of infection (MOI) of ∼1-5. Protein expression was carried out for ∼96 hours at 27 °C. Following expression, the pH of cell culture supernatant was adjusted to 8.0 by addition of 1 M Tris base, harvested, clarified, and concentrated using Centramate tangential flow filtration system.^52^ Alternatively, without prior concentration, Strep-tag II fused Fab proteins were purified directly from the pH-adjusted cell culture supernatant using the WET FRED system (IBA Lifesciences, Cat. No. 2-0910-001), which enables gravity-flow affinity purification. The concentrated supernatant was applied to a pre-equilibrated 4 mL bed volume Strep-Tactin 4Flow high-capacity resin (IBA Lifesciences, Cat. No. 2-1250-010) equilibrated in TBS (20 mM Tris-HCl pH 7.5, 150 mM NaCl) buffer. Fab bound resin was washed with six column volumes of TBS buffer, and Fab was eluted with three column volumes of TBS containing 7 mM desthiobiotin pH 7.5 (Neuromics, Cat. No. 2-1000-005). Eluted fractions were further purified by size-exclusion chromatography (SEC) using a Superose 6 10/300 GL column equilibrated in TBS buffer, concentrated to ∼2-5 mg/mL, and stored at −80 °C for further use. The C-term 10 aa construct yielded ∼2-2.5 mg per liter of culture, whereas cysteine-engineered variants exhibited lower yields of ∼0.7-1.5 mg per liter. Due to the presence of solvent-exposed thiol groups, cysteine-engineered Fabs were prone to intermolecular disulfide-mediated dimerization. To mitigate this, tris(2-carboxyethyl)phosphine (TCEP) (Sigma-Aldrich, Cat. No. C4706) was included at a final concentration of ∼1-2 mM to maintain reducing conditions, thereby preventing intermolecular disulfide formation and preserving monomeric Fab species. For native receptor purifications, C-term 10 aa and C-term 4 aa Fabs were used directly for AuNP conjugation without tag removal. For H1-2Cys, H1-3Cys, H4K-2Cys, and L1-2Cys variants, affinity tags were removed by in-house HRV 3C protease-His tag protein cleavage (1:50 molar ratio of protease and Fab, ∼2 mg/mL Fab, overnight on ice). Cleaved Fab was isolated by metal affinity chromatography to remove residual His-tagged protease and followed by Strep-Tactin affinity purification to remove Strep-tagged components. Affinity tag-free Fab samples were subsequently used for AuNP conjugation (**Supplementary Fig. 1a-b, 2**).

### Preparation of 15F1 Fab for crystallization

SEC purified C-terminal 10 aa Fab was subjected to affinity tag removal as described above. The resulting Fab was further purified by anion-exchange chromatography using a 5 ml HiTrap Q column (Cytiva, Cat. No. 17115301) equilibrated in buffer A (20 mM Tris pH 8.0, 10 mM NaCl, 0.1 mM EDTA, 0.1 mM methionine) and eluted with a linear gradient to buffer B containing 500 mM NaCl. Peak fractions were pooled and concentrated. For crystallization, the purified Fab was concentrated to 11.6 mg/mL and incubated at 20 °C. Using a 2 µL drop in hanging drop format, Fab was mixed at a 1:1 ratio with precipitant containing 0.1 M Tris (pH 7.0), 24% PEG 3350, and 20% ethylene glycol. Crystals were harvested and directly vitrified in liquid nitrogen without further cryopreservation.

### X-ray diffraction and data refinement

Shutterless X-ray diffraction data were collected at the Advanced Light Source beamline 8.2.1. The data were processed in XDS^53^ and Pointless/Aimless.^54^ The space group is P212121 with 4 molecules in the asymmetric unit. Analysis by Phenix Xtriage^55^ determined no evidence of twinning, however significant tNCS (24.4%) was seen. Initial phases were solved by molecular replacement with an AlphaFold model from the sequence including both heavy and light chains. Structures were then subjected to iterative model building and refinement using COOT^56^ and PHENIX^55^ software packages. Geometric and stereochemical validation was performed using MolProbity^57^ in PHENIX. Data collection and refinement statistics are provided in Supplementary Table 1.

### Fab reduction, test AuNP-Fab conjugation and PEGylated conjugate preparation

TBE buffer was prepared as a 10x stock solution containing 1 M Tris, 1 M boric acid, and 20 mM EDTA (pH 8.3). The stock was diluted to 1x working concentration and used for conjugation reactions, native gel electrophoresis, and SEC and FSEC. TCEP was prepared as a 100 mM stock solution in 100 mM Tris buffer (pH 8.0), aliquoted, and stored at −20 °C until use. Native AuNP gels (12% acrylamide/bis-acrylamide, 10% glycerol, 0.5x TBE) were prepared by standard polymerization using ammonium persulfate (APS; 0.05% w/v) and TEMED (0.001% v/v). Following polymerization, gels were maintained in a hydrated state and stored at 4 °C by wrapping in moist tissue paper until use.

Fabs samples at ∼2 mg/mL were incubated with 2-3.5 mM TCEP in TBE buffer (100 mM Tris, 100 mM boric acid, 2mM EDTA, pH 8.3) for 60 minutes at 37 °C in a water bath. Reduced samples were clarified by ultracentrifugation (86,500 × g, 15 minutes) and immediately subjected to SEC equilibrated in TBE buffer to remove reductant. Alternatively, TCEP was removed using Zeba desalting columns (ThermoFisher, Cat. No. 89890). The 15F1 Fab remained stable across 0.5-60 mM TCEP, with optimal stability up to 20 mM (**Supplementary Fig. 3**). For experiments comparing conjugation efficiency between different 15F1 Fab cysteine mutants, all Fabs were reduced with 2.5 mM TCEP at 37 °C for 60 minutes.

For large-scale Fab-AuNP conjugate preps, test scale conjugation reactions were first performed to optimize Fab to AuNP ratios using 3-MBA coated AuNPs. Reactions were assembled in 1x TBE buffer with varying Fab and AuNP volumes (6:1, 4:1, 1:1, 1:2, 1:4, 1:6, and 0:1) and incubated at 37 °C for 30 minutes. Following incubation, loading buffer (20% glycerol in TBE) was added, and samples were analyzed by native PAGE. Optimal conjugation conditions were defined as those producing the highest relative intensity of the 1 Fab:1 AuNP conjugate band. In experiments comparing conjugation efficiency across Fab cysteine mutants, a 2:1 ratio was used. Large-scale conjugation reactions were performed under optimized conditions and resolved on 12% acrylamide, 10% glycerol native gels (**Supplementary Fig. 4a**) in TBE buffer under cold conditions. The band corresponding to the 1 Fab:1 AuNP conjugate was excised, cut into small pieces, and eluted in 5 mL TBE buffer overnight at 4 °C. Gel pieces were subsequently extracted with an additional 5 mL TBE buffer until clear. Eluted fractions were filtered through a 0.22 µm Steriflip vacuum filtration system (Fisher, Cat. No. SCGP00525), pooled, and concentrated. mPEG-SH, Mw 550 (Creative PEGworks, Cat. No. PLS-607) was then added to final concentration of 0.5 mM, followed by incubation at 37 °C for 30-60 minutes with gentle mixing. PEGylated conjugates were further purified by FSEC using a Superose 6 column equilibrated in TBE buffer, monitoring AuNP absorbance (∼500 nm) and tryptophan fluorescence (Ex: 280 nm; Em: 335 nm). Fractions exhibiting co-elution of AuNP absorbance and tryptophan fluorescence were collected, concentrated, and stored at 4 °C (**Supplementary Fig. 4b**). The Fab-AuNP conjugates remained stable for approximately 3-6 months at 4 °C. The receptor-binding ability of the prepared conjugates was assessed using recombinant rat GluA2 receptor-GFP at various molar ratios. Complex formation was monitored by co-elution in the GFP fluorescence and 500 nm absorbance channels (**Supplementary Fig. 4c**). The concentration of Fab-AuNP conjugates was estimated by SDS-PAGE, comparing the bands from a Fab-AuNP sample and to those from unconjugated Fab standards at known concentrations.

### Stability of 15F1 C-terminal 10 aa Fab-AuNP conjugates under reducing conditions

PEGylated Fab–AuNP conjugates were incubated with reduced L-glutathione (0.2, 2.5, 10, 30, or 100 mM) at 37 °C for 60 minutes. Samples were centrifuged at 1,500 × g for 10 minutes and analyzed by FSEC using a Superose 6 Increase 10/300 GL column equilibrated in TBE buffer.

### Preparation of native AMPA receptor –Fab-AuNP conjugate complexes

Purification of native mouse AMPAR using the anti-GluA2 15F1 Fab-AuNP was done as previously described for the C-term 10 aa and C-term 4 aa conjugates.^27^ For the C-term 10 aa conjugate prep, the hippocampi were dissected from 10 male and 4 female adult vGluT1-mScarlet mice (total tissue mass = 675 mg). For the C-term 4 aa conjugate prep, the whole brain was dissected from 2 female adult C57BL/6 mice. Dissected tissue was resuspended in 5-7.5 mL of homogenization buffer (20 mM Tris-HCl pH 8.0, 150 mM NaCl, 0.8 μM aprotinin, 2 μg/mL leupeptin, 2 μM pepstatin A, 2 μM ZK-200775, 2 μM JNJ-55511118, 50 μM (R,R)-2b) per gram of tissue on ice. A Dounce homogenizer was used to homogenize the tissue, and the homogenized sample was diluted 1:1 with solubilization buffer (homogenization buffer plus 4% w/v digitonin). The solubilized tissue was nutated for 15 minutes at 4 °C then centrifuged at 4000 *g* for 3 minutes at 4 °C and run through a 0.22 μm filter.

The 15F1 Fab-AuNP conjugate, containing a C-terminal Strep-tag, was added to a concentration of approximately 40 nM and the sample was nutated at 4 °C for 15 minutes, then passed over StrepTactin Superflow or XT 4Flow resin equilibrated in purification buffer (20 mM Tris-HCl pH 8.0, 150 mM NaCl, 2 μM ZK-200775, 2 μM JNJ-55511118, 1 μM (R,R)-2b, 0.075% w/v digitonin) by gravity flow. Native AMPARs bound to 15F1 Fab-AuNP were eluted with purification buffer augmented with 5 mM desthiobiotin or 1X BXT buffer (iba, Cat. No. 2-1042-025), concentrated to 120 µL, and further purified over a Superose 6 Increase 10/300 GL size exclusion column, using an HPLC. The tryptophan fluorescence at excitation/emission wavelengths of 280/335 nm and AuNP absorbance at 500 nm were monitored and peak fractions corresponding to the 15F1 Fab-AuNP bound native AMPAR species were collected and concentrated to 20 μL.

For all other 15F1 Fab-AuNP conjugates (H1-2Cys, H1-3Cys, L1-2Cys, and H4K-2Cys), the C-terminal Strep-tag was cleaved prior to preparation with native AMPA receptor, to ensure the affinity tag was not affecting the observed AuNP localization. Whole brain tissue from male and female adult PSD95-GFP^27^ or C57BL/6 mice was dissected, homogenized, solubilized, and centrifuged as described above, using 2-3 mice per prep. An anti-GluA1 single-chain variable fragment fused to GFP and a Strep tag, 11B8 scFv-GFP-Str,^58^ was added to the filtered solubilized supernatant to a final concentration of 40 nM and the sample was nutated at 4 °C for 30 minutes. The solution was then passed two times over 2 mL of Strep-TactinXT 4Flow high-capacity resin equilibrated in purification buffer by gravity flow. The resin was washed with 20 mL of purification buffer and eluted with 10 mL of purification buffer + 0.002 mg/mL 3C protease (Takara), and the elution was concentrated to 120 μL. The 15F1 Fab-AuNP was added to the elution sample at an approximate concentration of 100-200 nM and incubated for 1 hour at 4 °C before purification over a Superose 6 increase column by HPLC, as described above.

The concentrated samples were diluted 1:1-1:8 in purification buffer and 3 μL was applied to Quantifoil R2/1 Cu or Au 200 or 300 mesh grids with a 2 nm continuous carbon film that had been glow discharged for 30 seconds at 15 mA with a PELCO easiGlow system. The grids were blotted and plunge frozen into a 35/65% ethane/propane mix using an FEI Vitrobot set to 4 °C, 100% humidity, 30 seconds wait time, 2.5-3.5 seconds blot time, and 0 blot force. Grids were screened on an FEI Glacios to assess ideal samples for data collection that contained relatively low concentration, well-spaced particles, aiming to minimize particle aggregation and the overlap of nearby particles within extracted boxes during processing.

### Cryo-EM data collection and processing of native AMPAR –15F1 Fab-AuNP complexes

Single particle cryo-EM grids with mouse native AMPAR bound to anti-GluA2 15F1 Fab-AuNP conjugates were collected on 300 kV Thermo Fisher Krios microscopes with either a Gatan K3 camera and GIF BioQuantum energy filter set at a slit width of 8 eV or a Thermo Scientific Falcon4i and Selectris X, and for some samples with a spherical aberration corrector (C-term 10 aa, C-term 4 aa, and H1-2Cys). Data were collected using a multishot pattern in SerialEM at a magnification of 64,000-165,000x (pixel size of 0.7252-1.088 Å) and defocus range of −0.5 to −1.5 or −0.5 to −3.5 µm, with 50 frames per movie at a total dose of 50 e/Å^2^. In total, 4,649-14,211 movies were collected for each sample (**Supplementary Table 3**).

Single particle cryo-EM datasets were processed in CryoSPARC v4.7.1^59^ using an identical strategy as described below (**Supplementary Fig. 7-8)**. Movies were patch motion and CTF corrected, and micrographs with a CTF fit estimated resolution lower than 5 Å were removed from analysis. Particles were first selected by blob picking using a circular blob and particle diameter of 180-250 Å. Iterative rounds of 2D classification without 2D class re-centering, to limit centering of the dominant AuNP signal, resulted in classes with well aligned AuNP signal and adjacent blurred receptor density. A final round of 2D classification with a circular mask diameter of 150 Å resulted in a subset of classes now with well-defined receptor density and masked-out AuNPs. These classes were used for template-based particle picking, followed by iterative rounds of 2D classification and particle cleanup. The best particles out of 2D classification in the blob-picked and template-picked particle sets were combined and used for *ab initio* reconstruction and heterogeneous refinement into four classes.

Classes that contained clear receptor and AuNP density were selected and the set of 2:1 15F1 Fab-AuNP:AMPAR particles (or 1:1 15F1 Fab-AuNP:AMPAR for the L1-2Cys conjugate) were used for non-uniform refinement, resulting in a map whose alignment was generally dominated by the AuNP signal. Particle subtraction using a mask around the AuNPs was used to remove the AuNP signal, followed by *ab initio* reconstruction and heterogeneous refinement into two classes. This produced one class in which some AuNP signal remained, resulting in a streaky blob density likely resulting from nearby AuNPs in the particle box, and one class in which AuNP signal was fully subtracted, resulting in a clear and well-aligned 15F1 Fab-bound AMPAR density. Non-uniform refinement followed by reconstruction without alignment using the original unsubtracted particles produced a map with clearly visible receptor and AuNP densities. Further local refinement of the AMPAR ATD:15F1 Fab region employing C2 symmetry, followed by symmetry expansion and refinement of a single ATD dimer bound to a 15F1 Fab, resulted in incrementally higher resolution reconstructions of the ATD-bound 15F1 Fab-AuNP. All refinements were carried out using AuNP signal-subtracted particles, then reconstructed without alignment using the original unsubtracted particles. For the H1-3Cys dataset, which contained the most particles, local CTF and motion correction was applied and further improved the resolution of the ATD_dimer_:15F1 Fab reconstruction.

Models of the AMPA receptor ATD from the native GluA2-containing mouse hippocampal AMPA receptor structure (PDBID 7LDD), the 15F1 Fab crystal structure from this study, and an AuNP:3-MBA structure^30^ were docked into the EM densities using the ‘fitmap’ command in ChimeraX^60^ for visualization and distance calculations. The ChimeraX marker placement tool was used to place a marker at the center of the AuNP densities within EM maps, contouring the maps to approximately the size of an AuNP for marker placement, for distance measurements.

### Acute mouse hippocampus slice preparation

Adult male and female vGlut1-mScarlet (HET)/PSD95-EGFP (HET) mice (6w-30w age) were used for the acute hippocampus slice preparation as described previously.^27^ Mice were anesthetized with isoflurane and brains were quickly removed after decapitation. Tissue blocks containing hippocampus were placed in a vibratome (Leica, VT1200). Horizontal brain slices (40 µm) were prepared at room temperature in one of two solutions. One solution was a HEPES-based buffer containing the following (in mM) 150 NaCl, 2.5 KCl, 2 CaCl_2_, 2 MgCl_2_, 10 HEPES, and 0.005 MK-801 (pH 7.3). A second solution was artificial cerebrospinal fluid (ACSF) buffer containing the following (in mM): 126 NaCl, 2.5 KCl, 1.2 MgCl_2_, 2.4 CaCl_2_, 1.2 NaH_2_PO_4_, 25 NaHCO_3_, 11 D-glucose, and 0.005 MK-801. The ACSF buffer was bubbled with 95%O_2_/5%CO_2_.

All animal procedures were performed in accordance with the guidelines of Oregon Health & Science University Institutional Animal Care and Use Committees (IACUC), consistent with the recommendations of the Panel on Euthanasia of the American Veterinary Medical Association (AVMA) and carried out by members of the EG laboratory approved by IACUC protocol IP00000905.

### Fab-AuNP immunolabeling and high pressure freezing of hippocampal slices

Hippocampal brain slices were incubated with corresponding the Fab-AuNP conjugate (2 µg/mL or ∼40 nM) in the HEPES buffer or 95%O_2_/5%CO_2_ oxygenated ACSF solution for 1 hour at room temperature on an orbital shaker (180 rpm). AMPA receptor antagonist, ZK-200775 (1 µM), and the positive allosteric modulator, (R,R)-2b (1 µM), were included throughout the experiments. For the experiment to examine the AMPA receptor desensitized state, AMPA receptor agonist, L-quisqualic acid (1 mM), was included throughout the experiments. After three washes of 15 minutess each, CA1 regions of the hippocampi were manually dissected with a scalpel knife and were then incubated in the HEPES buffer or in the 95%O_2_/5%CO_2_ oxygenated ACSF solution containing 20% dextran for 30 minutes prior to high-pressure freezing (HPF). In order to apply gold fiducials for tilt series alignment, PEGylated 10 nm gold fiducials (CP11-10-PA-1K-DI water-50, Nanoparts) were used. A volume of 100 µL of gold fiducials was spun down at 21,300 × g for 20 minutes and the supernatant was removed. After adding 100 µL of the HEPES or ACSF buffer, the centrifugation step was repeated. After removing the supernatant, 30-50 µl of HEPES or ASCSF buffer containing 20% dextran and 5% sucrose was added. This HEPES base cryoprotectant solution with gold fiducials was applied ∼5 to 10 seconds before HPF. Alternatively, CA1 slices were incubated in 95%O_2_/5%CO_2_ oxygenated ACSF containing 20% dextran with gold fiducials for approximately 30 minutes prior to HPF. In this case, oxygenated low Na ACSF solution containing the following (in mM): 75 NaCl, 2.5 KCl, 1.2 MgCl_2_, 2.4 CaCl_2_, 1.2 NaH_2_PO_4_, 25 NaHCO_3_, 11 D-glucose and 5% sucrose, was added immediately before HPF.

Prior to HPF, planchettes and grids were pretreated as described previously.^27^ The sample and 6 mm flat specimen planchettes were assembled at the loading station of a Leica EM ICE. Two CA1 tissue slices were placed on 200 mesh gold extra thick carbon film grids (CF200-Au-ET; EMS) with apical dendrites facing toward the center of the grid and CA1 cell layers were vertical, aligned with the rim mark located at the ‘12 o’clock’ position of the grid. This slice orientation enables us to target Schaffer collaterals, where CA3 axons synapse onto CA1 apical dendrites. After applying a small volume (∼1 µL) of sucrose containing cryoprotectant solution, a second planchette was placed on top of the sample grid with flat side down and the sample was immediately subject to HPF. After HPF, sample grids were clipped using a cryo-FIB autogrid ring and C-clip, with the carbon layer facing the autogrid ring side and the sample side facing the C-clip. The grid rim mark was oriented opposite the autogrid ring milling notch to preserve sample orientation for milling.

### Cryo-FIB milling of tissue samples

The HPF hippocampal brain slice grids were milled on an Aquilos 2 cryo-FIB/SEM microscope (FEI Company). After taking SEM tile images of an entire grid, sputter coating with a platinum layer (30 mA current for 30 seconds at 10 Pa pressure) and a gas injection system (GIS) deposition of a trimethyl(methylcyclopentadienyl) platinum layer (25 seconds) were applied before FIB milling. The grids were oriented with the Schaffer collaterals parallel to the ion beam path such that the resulting lamella targets could be chosen in 2-4 grid squares away from the center line on both left and right sides and up to 12 grid squares in a vertical orientation, depending upon the smoothness of the surface of the grid squares. The HPF brain tissue lamellae were made using the ‘waffle’ method^61,62^. At a 90° milling angle, two rectangular trench cuts were milled with an ion beam current of 15 nA until the front and back of the lamella target site were completely cleared. The sample shuttle was moved to a 20-40° milling angle, and the bottom section of the lamella site was removed. After the notch patterns were milled at a 20° milling angle on the sides of the lamella sites, another GIS deposition was applied for 20 seconds. Automated milling was performed using AutoTEM software with the milling angle of 20° and a set final lamella thickness of 140-160 nm. AutoTEM settings used for each of the steps were the following: rough milling at the ion beam current of 1 nA using rectangle patterns with a pattern offset of 1 µm, medium milling at 0.5 nA using cleaning cross sections (CCS) with a pattern offset of 0.8 µm, fine milling at 0.3 nA using CCS with a pattern offset of 0.6 µm, finer milling at 0.1 nA using CCS with a pattern offset of 0.4 µm, polishing 1 at 50 pA using CCS with over and under tilt of 0.5° and a pattern offset of 150 nm, and polishing 2 at 30 pA using CCS. After automated milling, the grids were tilted to +0.5° and −0.5° of the milling angle and manually polished with a cleaning cross section at 30 pA ion beam current to produce the final lamella. The milled grids were retrieved from the Aquilos 2 and stored in liquid nitrogen until they were imaged.

### Cryo-ET tilt series collection

Cryo-ET data on brain tissue lamella samples were collected on FEI Titan Krios 300 keV transmission electron microscopes equipped with either Thermo Scientific Falcon4i and Selectris X or Gatan K3 and BioContinuum HD detectors and energy filters, respectively. One microscope was also configured with a spherical aberration corrector. Tilt series were collected using a dose-symmetric tilt scheme with a 3° tilt step starting normal to the lamella at −20° and going to maximum tilts of −68° and 28° with SerialEM^63^ or Tomography 5 (Thermo Scientific). Movies were collected at pixel sizes of 1.9-2.3 Å, a defocus range of −2.5 to −7 µm, and a total exposure of 120-150 e^-^/A^2^. For the three different 15F1 Fab-AuNP conjugate immunolabeled brain tissue samples, 2,978 tilt series were collected across 30 different grids that contained a total of 84 lamella. Cryo-ET data on the single particle grid containing the purified H4K-2Cys 15F1 Fab-AuNP:AMPA receptor complex were collected using a collection scheme approximately equivalent to that used for the tissue lamella samples. A total of 186 tilt series were collected using a dose-symmetric tilt scheme with a 3° tilt step starting normal to the sample at 0° and going to maximum tilts of −48° and 48° with Tomography 5 (Thermo Scientific). Movies were collected at a pixel size of 2.4 Å, a defocus range of −3 to −5 µm, and a total exposure of 120 e^-^/A^2^.

### Tomogram reconstruction, segmentation, and annotation

For the brain tissue tomography data, all movies were motion corrected and binned to a pixel size of 2.5 Å using MotionCor3.^64^ Tilt series were initially reconstructed using AreTomo,^65^ and both motion-corrected tilt series and AreTomo reconstructions were individually reviewed to identify good quality tomograms that contain clear synapses. Selected tilt series were then aligned in Etomo by fiducial tracking, if sufficient 10 nm gold fiducials were present, or by patch tracking. CTF parameters were estimated using Ctfplotter^66^ and tomograms were reconstructed in Etomo by regular weighted backprojection with dose-weighting and 3D CTF correction. Tomograms were binned by 4 to a pixel size of 1 nm. Even and odd frame split versions of each tomogram were prepared in the same way and used for denoising model training.

Reconstructed tomograms were denoised using DeepDeWedge,^67^ which was trained on even and odd frame split versions of 2 to 3 tomograms, and training independent models for data collected on different detectors. Membranes were segmented from the DeepDeWedge denoised tomograms with MemBrain-seg,^68^ using the pretrained MemBrain_seg_v10_alpha.ckpt model checkpoint. Segmented membranes were imported into Blender using the MolecularNodes plugin,^69^ and the segmentations were manually thresholded and cleaned to separate cellular compartments and remove noise. Segmented membranes corresponding to individual presynaptic compartments, presynaptic vesicles, and postsynaptic compartments were manually selected and saved as independent mesh files for further analyses. To visualize positions of AuNPs between pre- and postsynaptic compartments, we calculated a minimum intensity projection (MIP) of a subvolume centered around representative AuNPs along a vector from the presynaptic to the postsynaptic compartment (**Supplementary Fig. 11a-b**).^70^

### Tomogram AuNP picking

A template for AuNP detection was generated from three tomograms of purified C-term 10 aa 15F1 Fab-AuNP conjugate, collected with identical imaging parameters as tissue tomograms (voxel size 1 nm, equivalent tilt scheme and defocus range). AuNPs in these tomograms were identified using the findbeads3d program in IMOD,^66^ yielding 28,409 detections. Subtomograms of 12 pixels were extracted at each detection and averaged to generate a template that captured characteristic aberrations present in the tomographic data.

Template matching was performed using pytom_match_pick^71^ with the averaged AuNP template, without rotational search and with spectral whitening. Initial candidate positions were identified as local maxima with a cross-correlation score above 0.3 and a minimum inter-peak distance of 2 pixels. To obtain sub-pixel accurate AuNP positions, subtomograms extracted at each candidate position were fit with a 3D Gaussian model. The center of the fitted Gaussian provided refined coordinates and the Gaussian amplitude served as an intensity metric for each AuNP feature. Neither the template matching score nor the Gaussian amplitude alone provided reliable discrimination of true AuNP detections, however, and optimal thresholds varied substantially between tomograms and synaptic regions due to differences in detector type, specimen thickness, and local alignment quality. Therefore, a range of template matching score and Gaussian amplitude thresholds were manually screened to maximize the accuracy of automated AuNP picking for each synaptic region.

Afterwards, all template-based picks were manually checked and remaining false positives and negatives were corrected to the best of our ability. In tomograms with visible platinum redeposition artifacts from cryo-FIB milling at the edges of the lamella, all putative AuNP picks within ∼6 nm of the tomogram edge were excluded from analysis. Additionally, AuNPs in tomogram subregions with locally poor alignment were excluded if they could not be confidently distinguished from noise or sufficiently improved by AuNP-guided local realignment (see below).

### AuNP-guided local tomogram realignment

When tomograms contained elongated AuNP features, suggestive of poor alignment of the tilt series or suboptimal motion correction, we carried out a local realignment that leveraged the AuNP features. The center points of regions of interest were manually identified and template-based AuNP picks or coordinates of darkest voxels within a radius of 120 nm of the center were used for alignment. A three-dimensional Gaussian model was fit to the density around each AuNP pick to determine sub-pixel precise centers and standard deviations (σx, σy, σz). Points above the 95th percentile in any dimension were filtered, and to distinguish genuine AuNPs from noise and gold fiducial markers, sigma values were constrained using thresholds corresponding to the estimated AuNP size, taking into account the missing wedge distortion (2 × 2 × 5 nm).

The refined AuNP centers were reprojected into the tilt series coordinate system using the alignment parameters generated by IMOD. A synthetic tilt series was generated by rendering a disk with a radius of 1.75 nm at each reprojected AuNP position. For each tilt angle, a cross-correlation map between the synthetic and experimental tilt images was calculated with a maximum search range of 25 nm. The position of optimal alignment was determined by fitting a two-dimensional Gaussian to the cross-correlation peak, and the resulting translational offsets were used to update the tilt series alignment parameters (**Supplementary Fig. 11c**). This pipeline was implemented in python via the scipy^72^ and teamtomo^73^ packages. Realigned tomograms were subsequently reprocessed through the previously described tomogram denoising, membrane segmentation, and annotation workflow.

### AuNP distance analyses

Presynaptic and postsynaptic cleft membranes within tomograms were defined as the regions of the segmentation meshes with vertices within 10-40 nm of each other at each synaptic region. Synaptic-localized AuNPs were retained for analysis by filtering AuNP picks to include only the subset within 30 nm of the postsynaptic cleft membrane. A k-dimensional tree built on all AuNP positions was used to calculate the distance to the nearest neighboring AuNP for each position. The edge of the outer and inner membrane segmentation at the cleft was classified based on the dot product of each vertex normal vector with the closest adjacent vertex normal on the opposing membrane (n_pre_ ·n_post_ < 0 defined the “outer” cleft-facing surface). The distance to the nearest vertex on the outer and inner mesh faces of the cleft membranes was calculated for each AuNP and the average of these distances was used to approximate the distance between each AuNP and the ‘center’ of the postsynaptic membrane.

### Subtomogram averaging of purified AMPA receptors

Motion correction and CTF estimation of tilt series collected on the purified H4K-2Cys 15F1 Fab-AuNP: AMPA receptor sample were done in WarpTools v2.0.0.^74^ Movies were binned to a 4.8 Å pixel size and tilt series stacks were aligned in Etomo^75^ by patch tracking. Tomograms were reconstructed with WarpTools at a pixel size of 1 nm and AuNP template matching was performed as described above. Particle positions were picked at the center of all isolated pairs of AuNPs (those with a pair distance of 5-9 nm and with no other AuNP picks within 20 nm), resulting in 5,975 putative 2:1 15F1 Fab-AuNP:AMPA receptor picks. Particles were exported with WarpTools for processing in RELION-5 v5.0.1^32^ at a 4 Å pixel size in a 128-pixel box as 2D particles. These particles were run through an initial round of 3D refinement in RELION, resulting in an AuNP-focused map with reasonably well-resolved 15F1 Fab-AuNP:AMPA receptor density.

Several attempts at 3D classification of this particle set resulted in less than 5% of particles classifying into “junk” classes. Therefore, no particles were excluded from analyses. The refinement was continued with a mask around the receptor that excluded the AuNP signal, resulting in a modest improvement to the receptor density. The particles were recentered on the receptor and aligned to the ∼2-fold axis for another round of refinement with C2 symmetry and using a smaller sampling range. Independent refinements masking either the ATD or LBD:TMD regions resulted in 15-20 Å reconstructions of the receptor complex (**Supplementary Fig. 10a-d**). Fidder^70^ was used to erase the AuNP signal from the CTF corrected micrographs and an additional round of focused refinements in RELION using these “AuNP-erased” tomograms resulted in marginal improvement to both the ATD- and LBD:TMD-focused reconstructions (**Supplementary Fig. 10e-f**). Final map resolution ranges were estimated with RESOLVE.^76^

## Data Availability

The coordinates for the 15F1 Fab crystal structure are deposited in the Protein Data Bank (PDB) under accession code 38DC. The cryo-EM maps of 15F1 Fab-AuNP: native AMPA receptor complexes have been deposited in the Electron Microscopy Data Bank (EMDB) under accession numbers EMD78575-78593 (see Supplementary Tables 3-4).

## Acknowledgements

We thank Rachel Courtney for assistance with manuscript preparation and Gouaux lab members for input on experiments and the manuscript. We acknowledge the generous support and use of the HHMI Janelia Cryo-EM facility for FIB milling on an Aquilos2, which was operated by Momoko Shiozaki and Nikki Jean, and data collection on Krios microscopes, which were operated by Drs. Rui Yan, Shixin Yang, and James Jung (HHMI Janelia). A portion of this research was supported by NIH grant R24GM154185 and performed at the Pacific Northwest Center for Cryo-EM (PNCC) with assistance from Janette Myers and Nancy Meyer. Electron microscopy grid screening was performed at the Multiscale Microscopy Core, a member of the OHSU University Shared Resource Cores RRID:SCR_009969. The research reported in this publication used computational infrastructure supported by the Office of Research Infrastructure Programs, Office of the Director, of the National Institutes of Health under Award Number S10OD034224. The content is solely the responsibility of the authors and does not necessarily represent the official views of the National Institutes of Health. E.G. is an investigator of the Howard Hughes Medical Institute and acknowledges generous support from Bernard and Jennifer LaCroute.

## Funding Statement

E.G. discloses support for the funding of this work from the NIH, grant number R01NS038631, and from the Howard Hughes Medical Institute. C.J.S. is supported by grant number CA253730 from the National Cancer Institute at the National Institutes of Health. J.N.P. discloses support for the funding of this work from the NIH, grant number R35GM142486.

## Author Contributions

G.D. and C.J.S. prepared Fab-AuNP conjugates. G.D. performed conjugation and stability assays. C.J.S. purified native receptors and performed single particle cryo-EM analyses. A.M. and C.J.S. prepared brain tissue cryo-ET samples. C.J.S. collected and processed cryo-ET data. J.E., and E.V.S. assisted with analysis of cryo-ET data. G.D. and J.N.P. prepared 15F1 Fab samples for crystallography studies and J.N. collected and processed crystallography data. E.G., C.J.S., and G.D. conceptualized the study and wrote the manuscript, with input from all authors.

## Competing Interest Statement

The authors declare no competing interests.

## Notes

### Competing Interest Statement

The authors have declared no competing interest.

