## Supplementary Information for "Programmable gold nanoparticle conjugates enable precise AMPA receptor localization at brain synapses by cryo-ET"

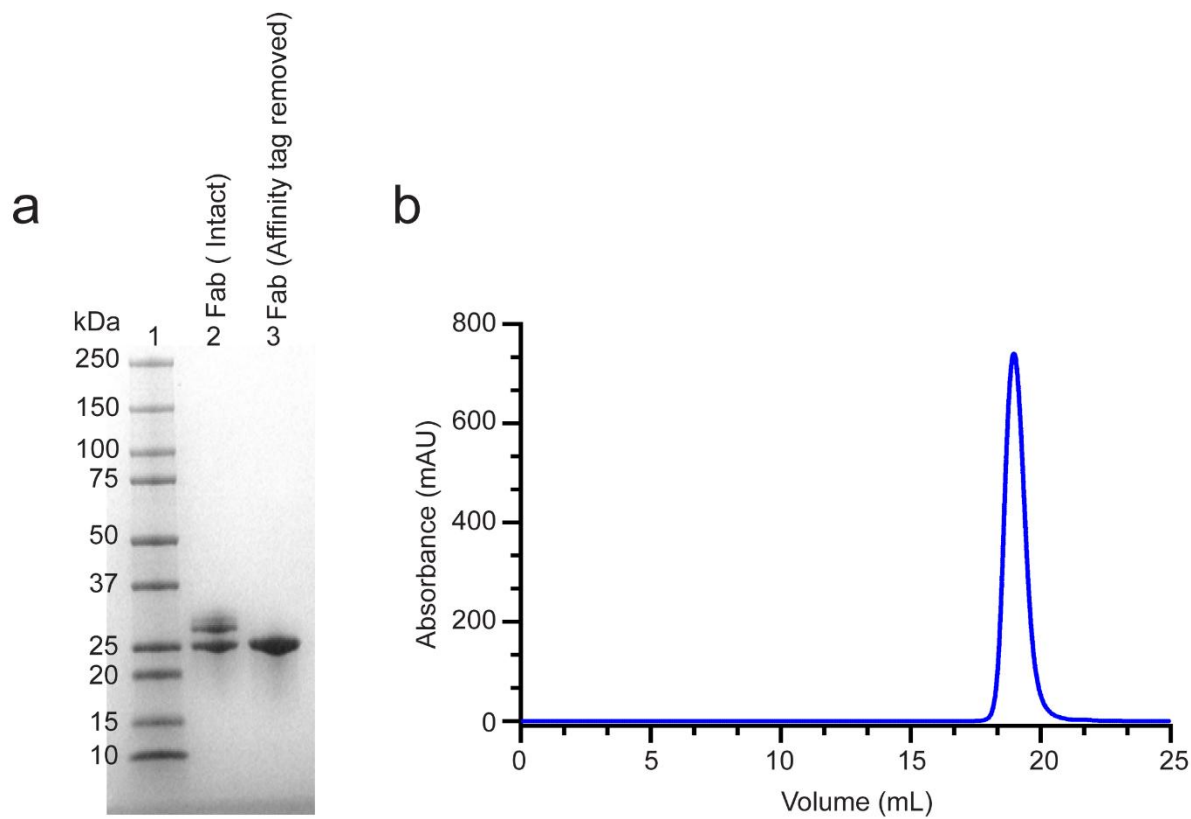

**Supplementary Figure 1: Preparation of wild type 15F1 Fab (C-term 10 aa) for crystallization.** **a)** Insect cell secreted Fab was purified by tandem Strep-Tactin affinity chromatography followed by SEC. The C-terminal Strep tag was removed by 3C protease digestion. The cleaved tag and residual uncleaved Fab were subsequently removed using reverse Strep affinity chromatography. Samples were analyzed by SDS-PAGE (12% TGX gel) under reducing conditions with 2-BME. Lane 1, Precision Plus Protein™ Dual Color Standards (Bio-Rad, Cat. No. 1610374); lane 2, intact Fab; lane 3, tag-removed Fab. **b)** 3C cleaved 15F1 Fab was further purified by anion exchange chromatography using a 5 ml HiTrap Q column. The purified sample, prior to crystallization, was evaluated by FSEC using intrinsic tryptophan fluorescence (Ex: 280 nm; Em: 335 nm) on Superose 6 Increase column equilibrated in TBS. The Fab eluted at 18.9 mL as a single, symmetric peak, consistent with monodisperse preparation.

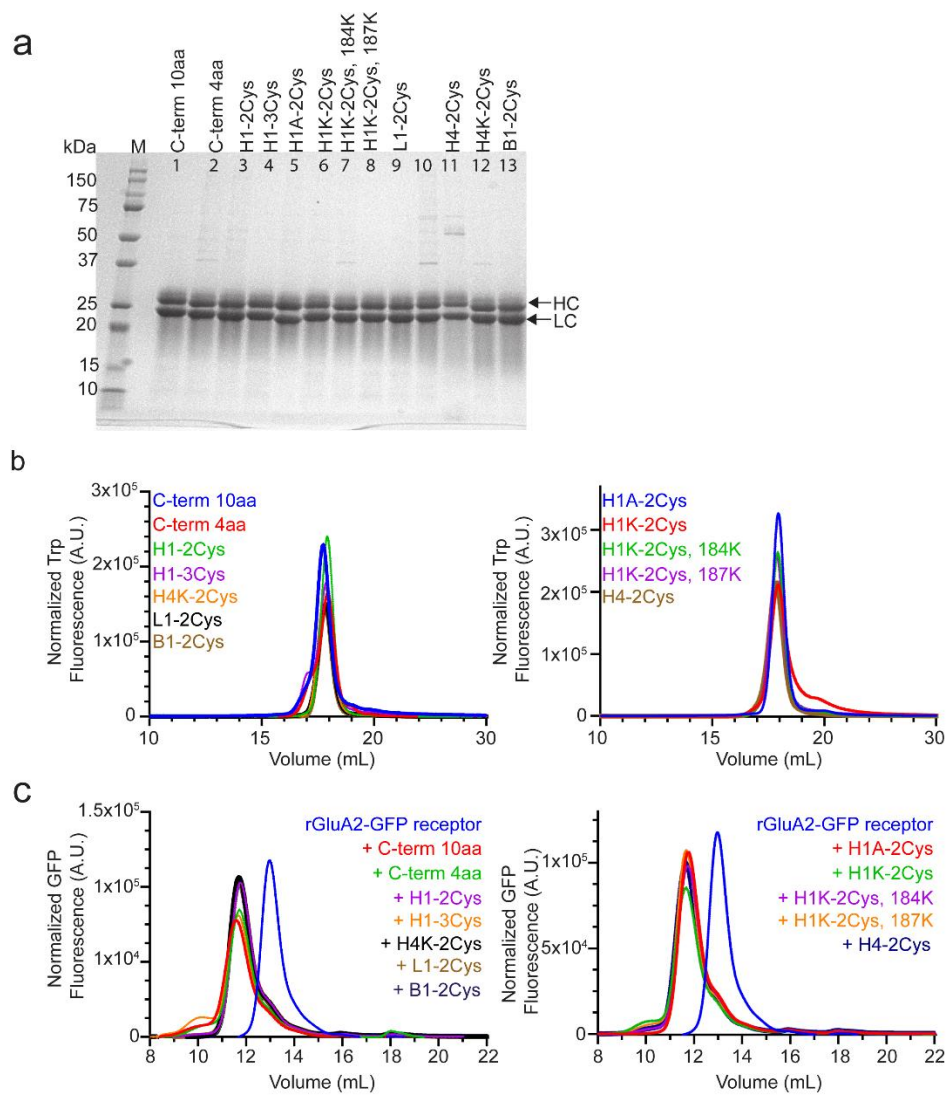

**Supplementary Figure 2: Biochemical and receptor binding characterization of 15F1 cysteine engineered Fab variants.** **a)** SDS-PAGE analysis of purified 15F1 Fab variants listed in Supplementary Table 2. All 12 mutants were expressed and purified, and their purity and integrity were assessed by SDS-PAGE (12% TGX gel) under reducing condition with 2-mercaptoethanol. Lane M, Precision Plus Protein™ Dual Color Standards (Bio-Rad, Cat. No. 1610374); Lanes 1–13 (except Lane 10) corresponds to individual Fab variants. Heavy chain and light chain bands are indicated by arrows. **b)** FSEC analysis of purified Fab variants using intrinsic tryptophan fluorescence (Ex: 280 nm/ Em: 335 nm). Samples were clarified by ultracentrifugation ($86,500 \times$

g, 20 min) prior to injection onto Superose 6 Increase column equilibrated in TBS. Left, Fab variants used for single-particle analysis; right, variants used for AuNP conjugation studies. c) FSEC peak shift analysis to assess binding of 15F1 Fab variants to recombinant GluA2 receptor-GFP. All Fab variants were incubated with receptor at ~1:2.5 molar ratio for 30 min on ice, followed by clarification by ultracentrifugation at $86,500 \times g$ for 20 min. Samples were analyzed by FSEC using GFP fluorescence (Ex: 480 nm/ Em: 510 nm). Analysis was performed on Superose 6 10/300 GL column equilibrated in buffer containing 20 mM Tris (pH 7.6), 150 mM NaCl, 1 mM DDM.

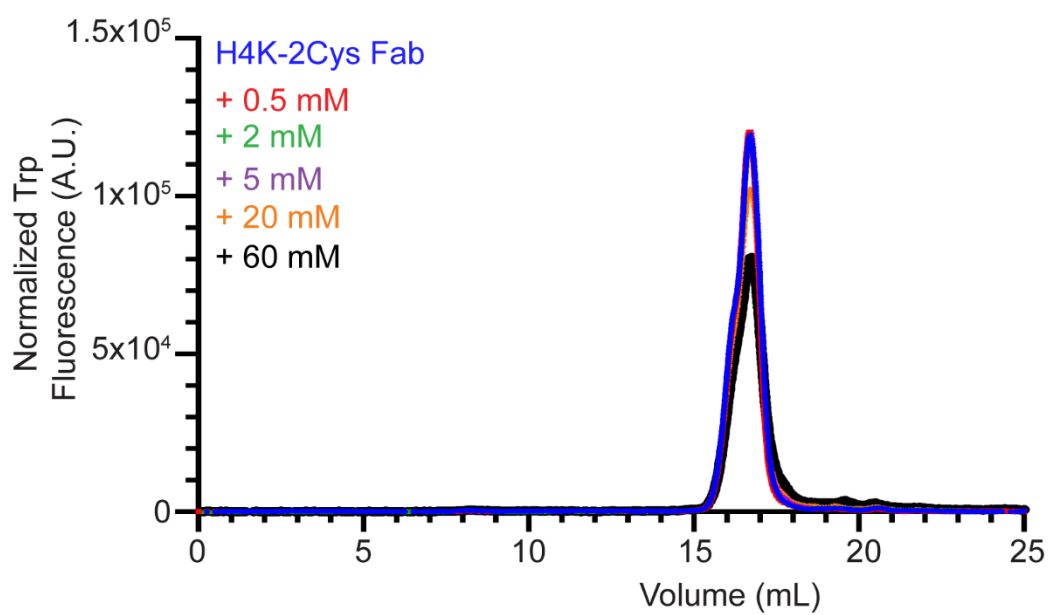

**Supplementary Figure 3: TCEP reduction stability analysis of 15F1 H4K-2Cys variant.** Fab samples were incubated with increasing concentrations of TCEP (0.5, 2, 5, 20, and 60 mM) for 1 h at 37 °C. Samples were subsequently clarified by centrifugation at $86,500 \times g$ for 20 mins, and supernatants were analyzed by FSEC using intrinsic tryptophan fluorescence (Ex 280 nm; Em 335 nm). Analyses were performed using Superose 6 10/300 GL column equilibrated in TBE buffer.

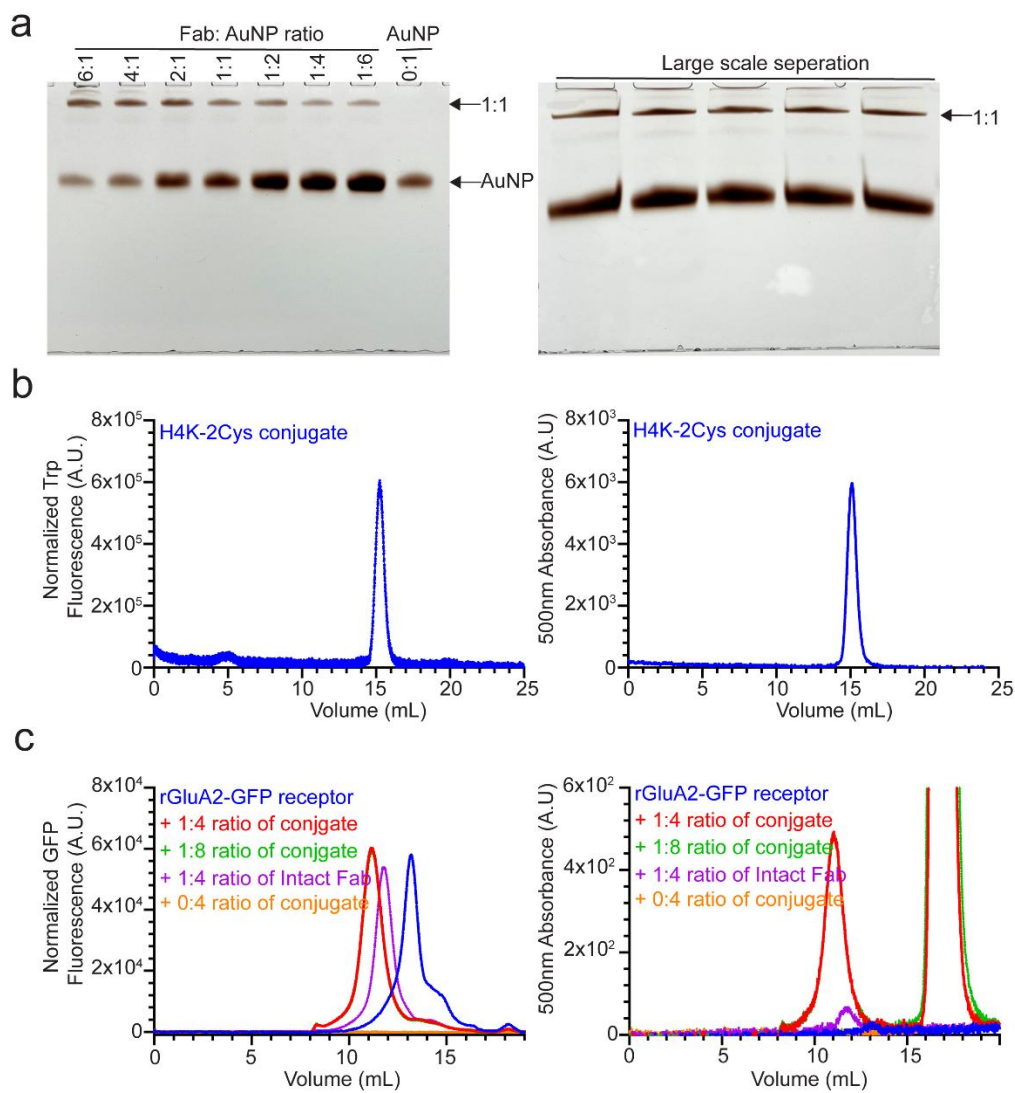

**Supplementary Figure 4: Preparation of affinity tag removed, PEGylated H4K-2Cys Fab-AuNP conjugates for native GluA2 AMPAR purification and synapse labeling.** **a)** Affinity tag removed H4K-2Cys Fab was conjugated to AuNP at varying Fab:AuNP volume ratios (6:1, 4:1, 1:1, 1:2, 1:4, 1:6, and 0:1) in TBE buffer at 37 °C for 30 min in total reaction volume of 15 μL. The initial concentrations were 2.45 mg/mL for Fab and ~10 mg/mL for AuNP. Conjugates were resolved using a 12% native AuNP gel (Left panel). A representative large-scale preparation is shown (Right panel), and 1:1 Fab-AuNP conjugate band was excised and recovered by gel elution using TBE buffer. **b)** Eluted conjugates were PEGylated with 0.5 mM mPEG-SH MW 550 and

further purified by FSEC to remove aggregates and displaced Fab or AuNPs. Fractions corresponding to 14.5-16.2 mL were collected and analyzed for quality assessment. The left panel shows tryptophan fluorescence, and the right panel shows 500 nm absorption channel. FSEC was performed using Superose 6 10/300 GL column equilibrated in TBE buffer. c) PEGylated Fab-AuNP conjugates were concentrated and incubated with rGluA2 AMPAR-GFP at varying molar ratios to assess complex formation. Binding was monitored by co-elution of GFP fluorescence and 500 nm absorption signals following incubation on ice for 30 min. Samples were analyzed by FSEC using a Superose 6 10/300 GL column equilibrated in buffer containing 1 mM DDM, 100 mM Tris, 100 mM Boric acid, 2.5 mM EDTA pH 8.3.

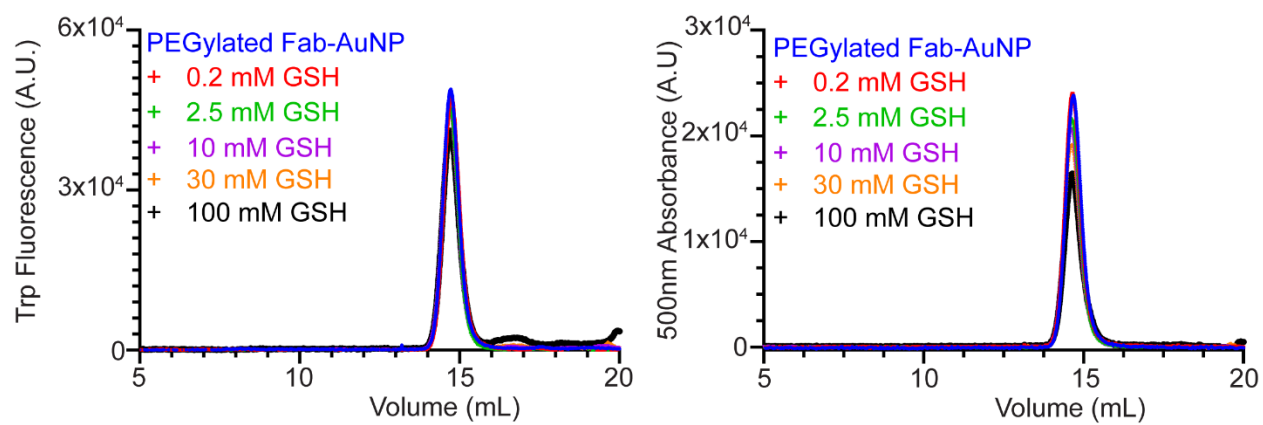

**Supplementary Figure 5: Stability of PEGylated H4K-2Cys Fab-AuNP conjugates under reducing conditions.** PEGylated H4K-2Cys Fab-AuNP conjugates were incubated with reduced L-glutathione (0.2, 2.5, 10, 30 and 100 mM) at 37 °C for 60 min. Following incubation, samples were centrifuged at $15,000 \times g$ for 15 min and analyzed by FSEC using a Superose 6 Increase 10/300 GL column equilibrated in TBE buffer. Elution profiles were used to assess conjugate stability under increasing reducing conditions.

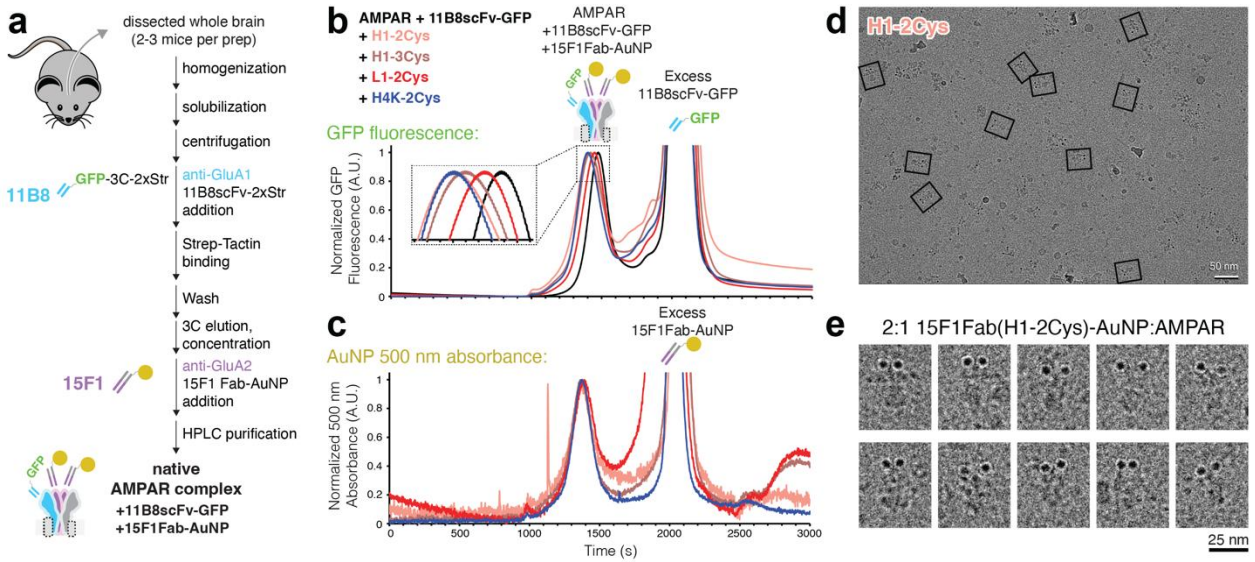

**Supplementary Figure 6: Small-scale affinity purification and cryo-EM of 15F1 Fab-AuNP: native AMPA receptor complexes.** **a)** Native AMPA receptor immunoaffinity pulldown from mouse brain tissue and 15F1 Fab-AuNP complex purification workflow. **b)** HPLC GFP fluorescence traces for native AMPA receptor:15F1 Fab-AuNP complex purifications. **c)** Equivalent HPLC 500 nm absorbance traces for samples from (b). **d)** Example screening micrograph from H1-2Cys 15F1 Fab-AuNP:native AMPA receptor complex sample. **e)** Particles from micrograph shown in (d).

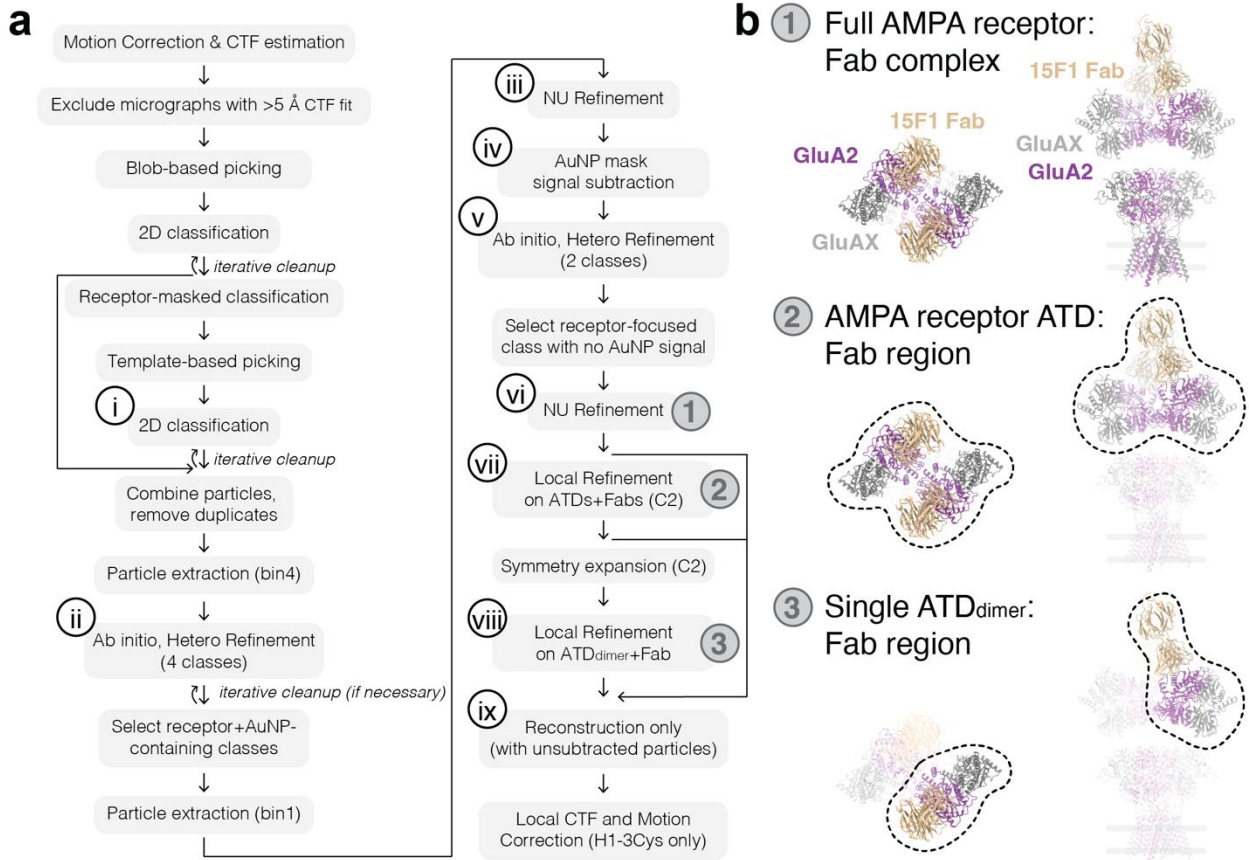

**Supplementary Figure 7: Single particle cryo-EM processing workflow for 15F1 Fab-AuNP: native AMPA receptor complexes – part I.** **a)** Processing steps for single particle cryo-EM analysis of all 15F1 Fab-AuNP:native AMPA receptor complex datasets. **b)** Illustration of the complex regions masked in focused refinements.

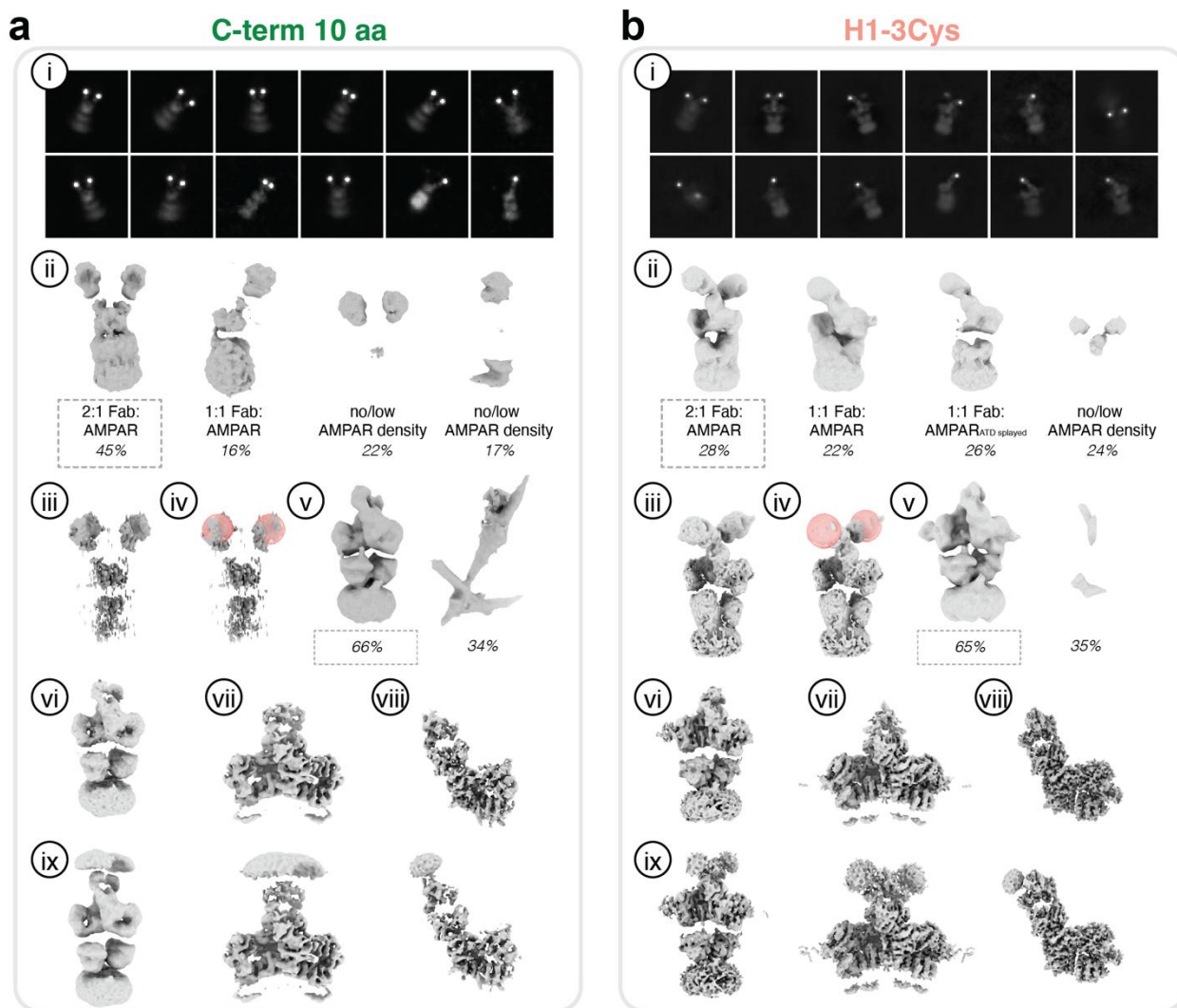

**Supplementary Figure 8: Single particle cryo-EM processing workflow for 15F1 Fab-AuNP: native AMPA receptor complexes – part II.** Example 2D classes (i) and cryo-EM maps (ii-ix) for processing steps shown in Supplementary Figure 7 for C-term 10 aa **(a)** and H1-3Cys **(b)** conjugates.

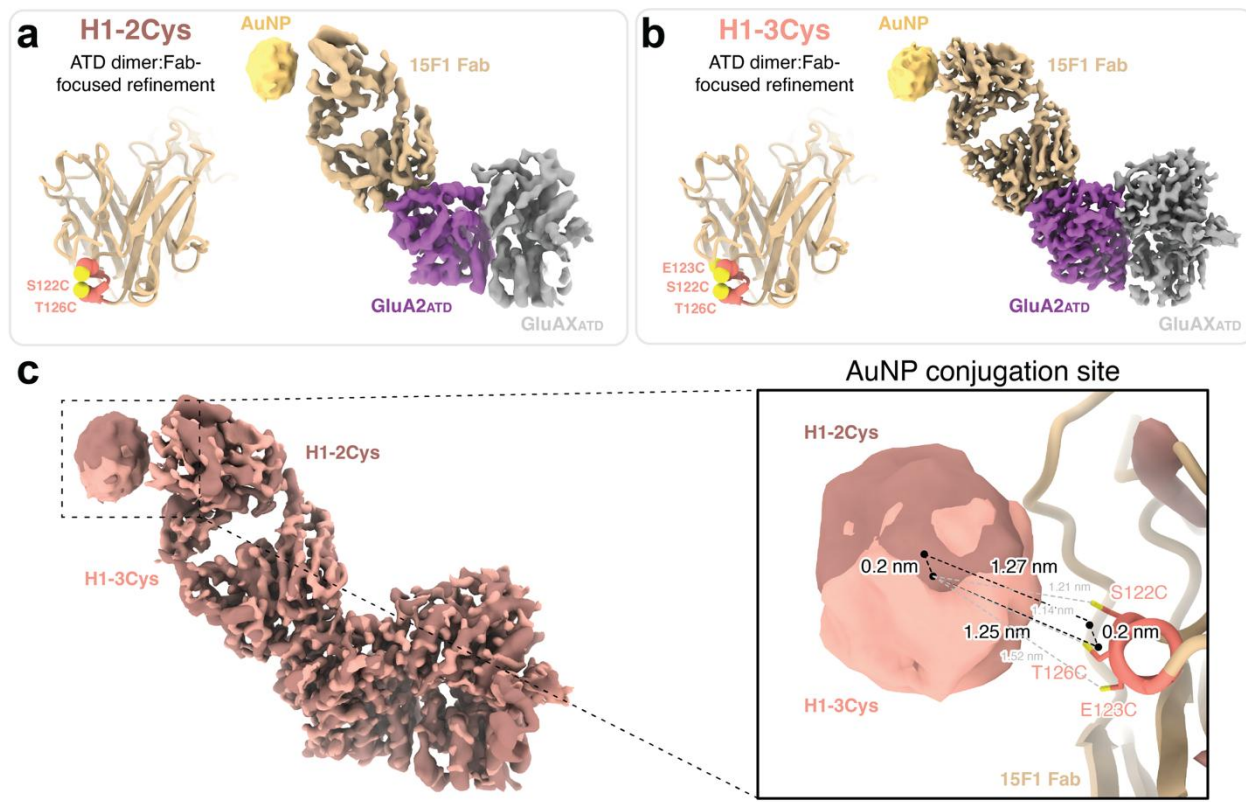

**Supplementary Figure 9: Comparison of AuNP conjugation in H1-2Cys and H1-3Cys conjugates.** Conjugation site residues and ATD<sub>dimer</sub>:Fab-focused refinement maps from single particle cryo-EM analysis of H1-2Cys (**a**) and H1-3Cys (**b**) conjugates. **c**) Alignment of H1-2Cys and H1-3Cys maps shown in (a-b) (left). Measurement of AuNP density center-to-center, AuNP density center-to-conjugation cysteine sulfurs centroid, and conjugation cysteine sulfurs centroid-to-centroid distances (right).

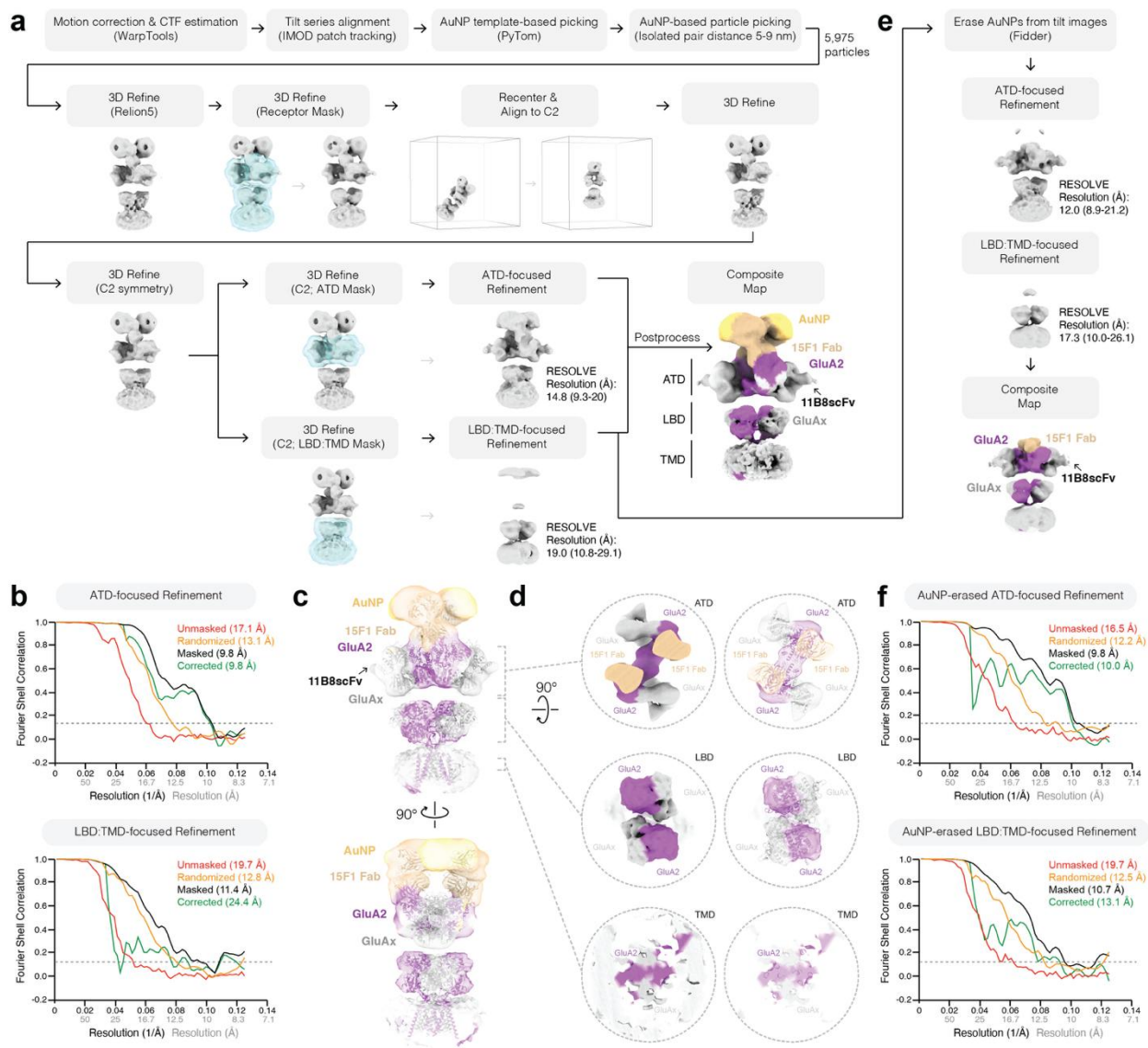

**Supplementary Figure 10: Subtomogram averaging workflow for purified H4K-2Cys 15F1**

**Fab-AuNP: native AMPA receptor complex. a)** Processing steps for subtomogram averaging

analysis of purified H4K-2Cys 15F1 Fab-AuNP: native AMPA receptor complex dataset. RESOLVE-estimated resolutions (median and range) are displayed next to final refinement steps.

**b)** Relion-5 Fourier shell correlation (FSC) analysis of ATD- and LBD:TMD-focused refinements from panel **a** with 0.143 cutoff denoted. **c)** Overlay of composite map with fit structure of native

GluA2-containing hippocampal AMPAR (PDB 7LDD). **d)** Clipped “top-down” views of subtomogram average map and aligned model fits for ATD, LBD, and TMD receptor regions. **e)** Subtomogram averaging analysis workflow for AuNP-erased data. **f)** FSC analysis of AuNP-erased ATD- and LBD:TMD-focused refinements from panel **e** with 0.143 cutoff denoted.

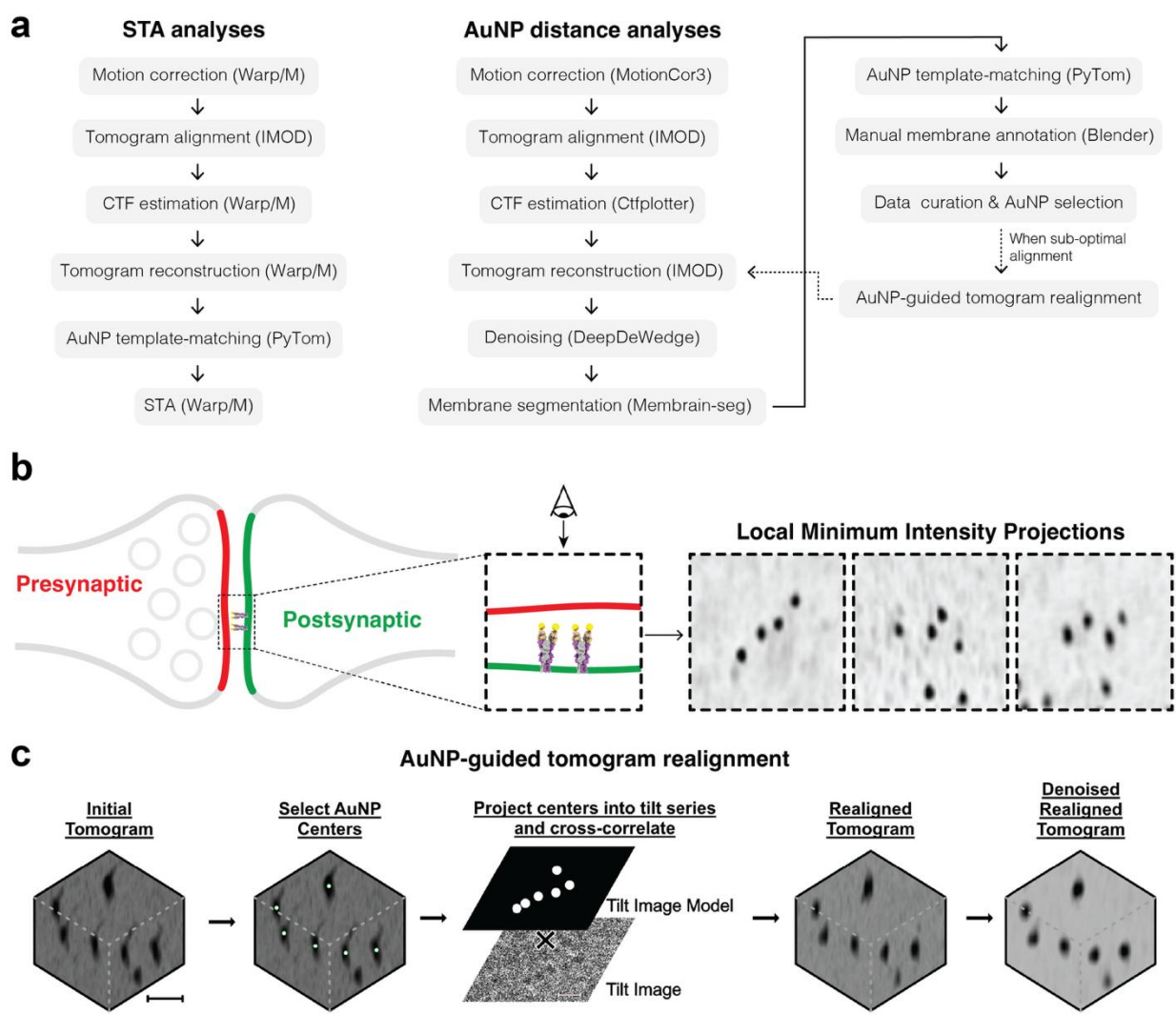

**Supplementary Figure 11: Cryo-ET data processing workflow.** **a)** Cryo-ET data processing steps in subtomogram averaging (STA) and AuNP distance analyses. **b)** Example subtomogram region displayed in top-down minimum intensity projection views of AuNP-labeled synapses. **c)** AuNP-based tomogram realignment workflow. Scale bars are 10 nm (Initial Tomogram) and 18.75 nm (Tilt Image).

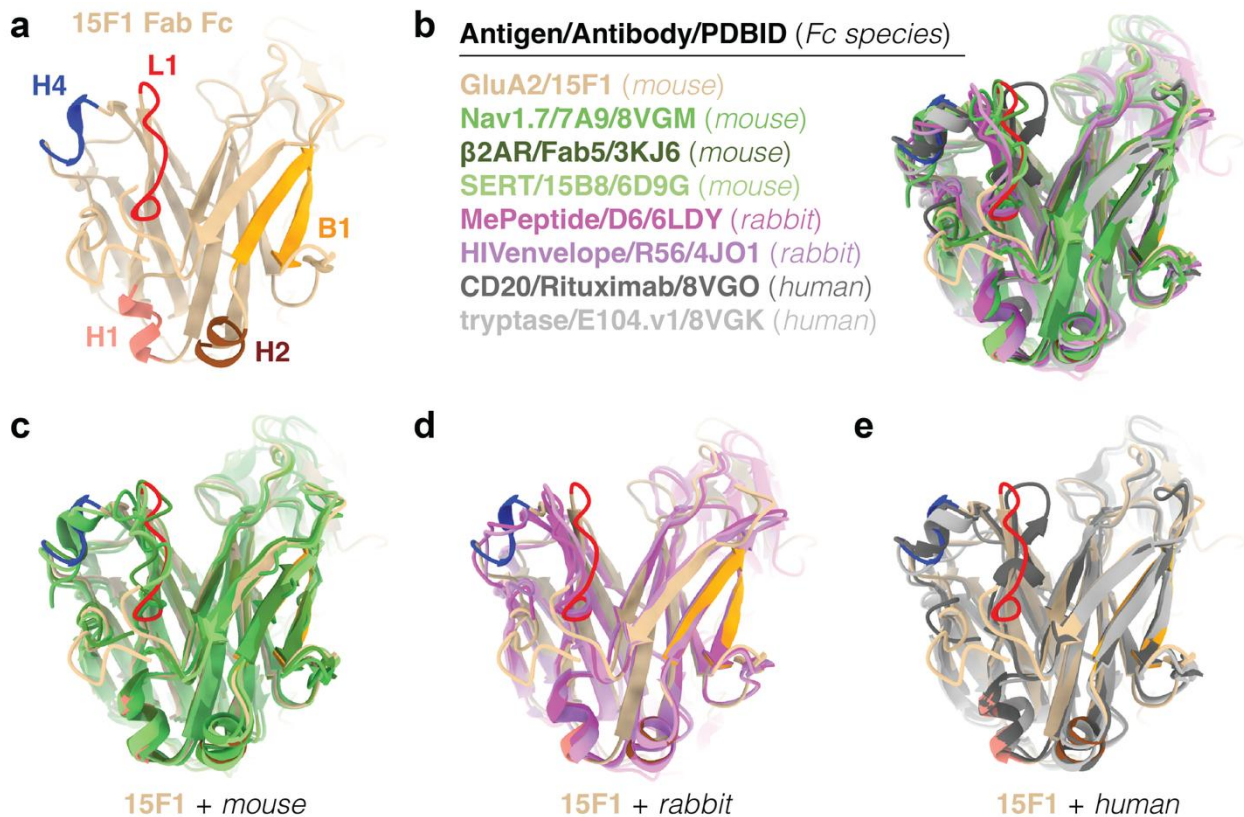

**Supplementary Figure 12: Structural comparison of Fab fragment constant domains across species.** **a)** Top-down view of 15F1 Fab constant domain with various structural elements denoted. **b)** Alignment of 15F1 Fab constant domain with mouse, rabbit, and human Fab fragment constant domain structures. Antigen, Antibody name, and PDB ID for each structure is noted. Alignment of 15F1 Fab constant domain with only the subset of mouse (**c**), rabbit (**d**), or human (**e**) Fab fragments shown in **b**.

**Supplementary Table 1: Crystallographic data collection and refinement statistics for 15F1**

**Fab.**

|  | 15F1 Fab<br>PDB: 38DC |
| --- | --- |
| **Data collection** |  |
| Space group | P212121 |
| Cell dimensions |  |
| *a*, *b*, *c* (Å) | 62.46, 64.28, 204.89 |
| $\alpha$, $\beta$, $\gamma$ (°) | 90, 90, 90 |
| Resolution (Å) | 46.81-2.50 (2.60-2.50)* |
| *R*<sub>merge</sub> | 0.061(0.468) |
| *I* / $\sigma I$ | 22.0(3.5) |
| Completeness (%) | 99.9(100.0) |
| Redundancy | 6.6(6.8) |
| **Refinement** |  |
| Resolution (Å) | 46.81-2.50 (2.60-2.50) |
| No. reflections | 27,9760(2,714) |
| *R*<sub>work</sub> / *R*<sub>free</sub> | 0.209/0.257 |
| No. atoms |  |
| Protein | 13252 |
| Ligand/ion | 0 |
| Water | 359 |
| *B*-factors | 48.94 (average) |
| Protein | 49.16 |
| Ligand/ion | N/A |
| Water | 40.36 |
| R.m.s. deviations |  |
| Bond lengths (Å) | 0.013 |
| Bond angles (°) | 1.45 |

\*Data collected on 1 crystal.

| **Label** | **Construct description** |
| --- | --- |
| C-term 10aa | LC 214C; HC 220C, 10 aa linker-3C-Twin Streptag (**IVPRDAGAKPC**) |
| C-term 4aa | LC 214C; HC 214C, 4 aa linker-3C-Twin Streptag (**IVPRDAGAKPC**) |
| H1-2Cys | LC **S**122C, **T**126C, **C**214S; HC **C**214S-3C-Twin Streptag |
| H1-3Cys | LC **S**122C, **E**123C, **T**126C, **C**214S; HC **C**214S-3C-Twin Streptag |
| H1A-2Cys | LC **S**122C, **E**123A, **T**126C, **C**214S; HC **C**214S-3C-Twin Streptag |
| H1K-2Cys | LC **S**122C, **E**123K, **T**126C, **C**214S; HC **C**214S-3C-Twin Streptag |
| H1K-2Cys, 184K2Cys | LC **S**122C, **E**123K, **T**126C, **D**184K, **C**214S; HC **C**214S-3C-Twin Streptag |
| H1K-2Cys, 187K2Cys | LC **S**122C, **E**123K, **T**126C, **E**187K, **C**214S; HC **C**214S-3C-Twin Streptag |
| B1-2Cys | LC **N**145C, **T**177C, **C**214S; HC **C**214S-3C-Twin Streptag |
| H4-2Cys | LC **C**214S; HC **S**192C, **S**196C, **C**220S-3C-Twin Streptag |
| H4K-2Cys | LC **C**214S; HC **S**193C, **S**197C, **E**198K, **C**220S-3C-Twin Streptag |
| L1-2Cys | LC **C**214S; HC **S**135C, **A**136C, **C**220S-3C-Twin Streptag |

**Supplementary Table 2:** List of 15F1 Fab constructs and corresponding mutation details. Surface exposed residues identified from 15F1 crystal structure were selectively mutated to cysteine to enable site-specific conjugation with 3-MBA coated AuNPs. Constructs span multiple structural regions (C terminus, H1, H4, B1 and L1) with cysteine positions in either heavy or light chain constant domains. For all except the C-term 10 aa and C-term 4 aa constructs, native terminal cysteines (HC C220 and LC C214) were substituted with serine to prevent undesired thiol reactivity. Mutated residues are indicated in bold within the construct descriptions. HC, heavy chain; LC, light chain; 3C, HRV 3C protease cleavage site; Twin-Strep tag, two Strep tags linked by 12 aa flexible linker.

**Supplementary Table 3: Single particle cryo-EM data collection and processing**

|  | C-term 10<br>aa | C-term 4<br>aa | H1-2Cys | H1-3Cys | L1-2Cys | H4K-2Cys |
| --- | --- | --- | --- | --- | --- | --- |
| **Data collection** |  |  |  |  |  |  |
| Magnification | 64k | 81k | 64k | 130k | 165k | 130k |
| Voltage (kV) | 300 | 300 | 300 | 300 | 300 | 300 |
| Electron exposure (e-/<br>Å<sup>2</sup>) | 50 | 50 | 50 | 50 | 50 | 50 |
| Defocus range (µm) | -0.5 to -1.5 | -0.5 to -3.5 | -0.5 to -<br>3.5 | -0.5 to -<br>3.5 | -0.5 to -<br>3.5 | -0.5 to -<br>3.5 |
| Pixel size (Å) | 1.070 | 1.061 | 1.088 | 0.9373 | 0.7252 | 0.9373 |
| **Processing** |  |  |  |  |  |  |
| **Full Receptor** | EMD-78586 | EMD-78587 | EMD-78588 | EMD-78589 | EMD-78590 | EMD-78591 |
| Symmetry imposed | C1 | C1 | C1 | C1 | C1 | C1 |
| Initial particle images<br>(no.) | 65,411 | 63,403 | 246,663 | 545,002 | 124,332 | 64,457 |
| Final particle images<br>(no.) | 19,625 | 34,471 | 38,591 | 71,433 | 26,403 | 19,983 |
| Map resolution (Å) | 10 | 9 | 10 | 7 | 12 | 9 |
| FSC threshold | 0.143 | 0.143 | 0.143 | 0.143 | 0.143 | 0.143 |
| **ATD:Fab(s)** | EMD-78580 | EMD-78581 | EMD-78582 | EMD-78583 | EMD-78584 | EMD-78585 |
| Symmetry imposed | C2 | C2 | C2 | C2 | C1 | C2 |
| Map resolution (Å) | 5.9 | 7.1 | 6.1 | 4.1 | 8.1 | 4.4 |
| FSC threshold | 0.143 | 0.143 | 0.143 | 0.143 | 0.143 | 0.143 |
| **ATD<sub>dimer</sub>:Fab** | EMD-78575 | EMD-78576 | EMD-78577 | EMD-78578 |  | EMD-78579 |
| Symmetry-expanded<br>particles (no.) | 39,250 | 68,942 | 77,182 | 142,866 | - | 39,966 |
| Map resolution (Å) | 4.2 | 7.0 | 4.6 | 3.5 | - | 4.2 |
| FSC threshold | 0.143 | 0.143 | 0.143 | 0.143 | - | 0.143 |

**Supplementary Table 4: Cryo-ET data collection and processing**

|  | C-term 10 aa, H1-2Cys,<br>or H4K-2Cys 15F1 Fab-AuNP labeled brain slice<br>lamella | H4K-2Cys 15F1<br>Fab-AuNP: native<br>AMPA (ATD-focused)<br>EMD-78592 | H4K-2Cys 15F1<br>Fab-AuNP: native<br>AMPA (LBD:<br>TMD-focused)<br>EMD-78593 |
| --- | --- | --- | --- |
| **Data collection** |  |  |  |
| Magnification | 42-53k | 53k | 53k |
| Voltage (kV) | 300 | 300 | 300 |
| Electron exposure (e-/Å<sup>2</sup>) | 120-150 | 120 | 120 |
| Defocus range (μm) | -2.5 to -7 | -3 to -5 | -3 to -5 |
| Pixel size (Å) | 1.9-2.3 | 2.4 | 2.4 |
| **Processing** |  |  |  |
| Symmetry imposed |  | C2 | C2 |
| Initial particle images<br>(no.) |  | 5975 | 5975 |
| Final particle images (no.) |  | 5975 | 5975 |
| Map resolution (Å) |  | 17.1 (Relion) | 19.7 (Relion) |
| FSC threshold |  | 0.143 | 0.143 |
| Resolution range (Å) |  | 9.3-20 (RESOLVE) | 10.8-29.1<br>(RESOLVE) |
